# Integrating multi-host modelling with empirical wildlife-livestock contacts reveals an essential population in a pathogen reservoir

**DOI:** 10.64898/2026.05.21.726803

**Authors:** Sébastien Lambert, Clara Meyers, Pauline Bouillot, Rémi Fay, Dominique Gauthier, Pascal Marchand, Ariane Payne, Elodie Petit, Anne Thebault, Timothée Vergne, Emmanuelle Gilot-Fromont

## Abstract

Infections at the animal-human or wildlife-livestock interfaces have severe health and socio-economic consequences. Combined with empirical data, mathematical models can contribute to a better understanding of the reservoirs of these infections, which is a priority for mitigating their impact by using appropriate management interventions. Taking brucellosis in the Bargy massif (French Alps) as an example of a zoonosis at the wildlife-livestock interface, we developed and calibrated a multi-host model integrating data on direct and environment-mediated cross-species contacts from field observations. Estimates of the basic reproduction number (*R*_0_) allowed to identify the population of Alpine ibex (*Capra ibex*) and its environment as an essential host in the reservoir, driving both pathogen maintenance (within-species *R*_0_≥1: 1.66, 95% credible interval: 1.42-2.03) and its transmission to livestock (between-species *R*_0_>0: 0.035, 0.01-0.05). Our approach can be adapted to other multi-host pathogens, which will contribute to improve the understanding and management of these complex systems.

## Introduction

Among known species of infectious agents in humans and domestic mammals, 58 to 90% are multi-host^1–3^, i.e. able to infect multiple host species. Multi-host pathogens are responsible for many of the major infectious diseases in humans, domestic animals and wildlife such as Ebola^4^, African swine fever^5^ of highly pathogenic avian influenza^6^, with severe consequences on human and animal health, economics, food security, and wildlife conservation^7–10^. They show complex epidemiological patterns, generally persisting in one or several host species or populations, but sometimes also affecting hosts in which they do not persist^11^. Consequently, in a specific eco-epidemiological setting, a reservoir of infection is defined as epidemiologically connected host populations or environments involved in two epidemiological functions^12,13^: (i) maintenance, i.e., the ability to maintain the pathogen within the ecosystem in the absence of other hosts, and (ii) transmission, i.e., the ability to transmit the pathogen to the population of concern or interest. Depending on their ability to contribute to one or both functions, and on whether they are essential to maintenance and/or to transmission, host populations in the ecosystem can play different roles^12–16^. Populations of the same species can even play different roles in different ecosystems, depending on the local context. For instance, the agent of bovine tuberculosis in the Iberian Peninsula is maintained either by wild boar (*Sus scrofa*), red deer (*Cervus elaphus*) or both, depending on the location^17^.

In practice, the identification of the host populations contributing to reservoirs of infection often starts by cumulating evidence from longitudinal epidemiological studies, analyses of prevalence and incidence patterns, experimental infections, pathogen genetics, and/or intervention studies^12,18–20^ to demonstrate both pathogen maintenance and transmission to the population of concern or interest. Examples where several lines of evidence were used to explore the potential role of multiple hosts include rabies in the Serengeti ecosystem in Tanzania^21^ and zoonotic schistosomiasis in sub-Saharan Africa^22^. However, the interpretation of empirical data may be difficult, owing to the complex and unique nature of multi-host reservoirs and their ecosystem. Many studies tend to conclude on the host population ability to maintain the pathogen based on high prevalence alone, but this is not sufficient^19^. Several mechanisms can lead to a high prevalence in a population despite not being able to maintain the pathogen on its own, such as high susceptibility to infection from other species but low shedding rates^19^. As an illustration, Eurasian blackbirds (*Turdus merula*) showed the highest Usutu virus prevalence in the Netherlands between 2016 and 2022, and yet were not able to maintain the virus on their own^23^.

The maintenance and transmission functions have been linked to a key epidemiological quantity, the basic reproduction number *R*_0_ (i.e., the average number of secondary infections resulting from one infectious individual in a totally susceptible population)^13,14,18^. For a pathogen to persist, the *R*_0_ must be higher than or equal to one, i.e., each infectious individual must, on average, infect at least one other individual. In multi-host systems, an overall *R*_0_ can be defined, which depends on the within-species *R*_0_ as well as the *R*_0_ between all possible pairs of species, i.e., the average number of secondary infections resulting from one infectious individual of one species in a totally susceptible population of another species^24,25^. These quantities can be estimated from empirical data using mechanistic models^14,17,26–28^, providing quantitative evidence to infer the maintenance and transmission potential of multiple hosts. By relying on clear assumptions about the main demographic and epidemiological processes of the system, mechanistic models can avoid biased claims based on empirical data alone, e.g., by distinguishing situations where high prevalence is associated with maintenance or not^19^. Despite these advantages, they remain largely underused for identifying the roles of different populations in a reservoir^19^, probably owing to the specific challenges posed by multi-host models^29^.

One of the main challenges of this type of model is the parametrisation of the multiple transmission rates^15^, which include both the rate of contacts appropriate for transmission and the probability that a contact between an infectious and a susceptible host does in fact lead to transmission^30^. In recent years, this challenge has been circumvented by using relative values of between-species transmission rates compared to within-species transmission rates to ease parametrisation^14,15^. While it is not usually possible to measure directly the absolute values of the transmission rates, it may be easier to obtain empirical evidence on the relative infectiousness, susceptibility and/or contact rates between hosts^15^. For instance, spatial overlap or proximity between host species or their habitats can be used as a proxy to inform relative contact rates^28,31^. However, to our knowledge, direct quantification of within and between-species contacts in natural settings have not yet been used to inform transmission rates in multi-host models, especially for diseases at the wildlife-livestock interface^32^.

The agents of brucellosis, mainly *Brucella abortus, B. melitensis*, and *B. suis*, are multi-host pathogens shared between wildlife, livestock, and humans^33^. Brucellosis is one of the most common zoonotic diseases worldwide, with an annual global incidence in humans estimated between 500,000^34^ and more than two million^35^ new cases. The primary hosts are domestic ruminants (cattle, sheep and goats) for *B. abortus* and *B. melitensis* and pigs for *B. suis*, with possible spillover to humans and wildlife. However, in some situations, maintenance in wild populations with spillover to livestock threatens decades of eradication programs in domestic ruminants, with human and animal health as well as economic consequences. This is the case for *B. abortus* in the Greater Yellowstone Ecosystem in the United States of America (USA)^36^ and for *B. melitensis* in the Bargy massif ecosystem in the French Alps^37^.

In this study, we combined for the first time a multi-host mechanistic model of *B. melitensis* transmission with observational data, including field observations of contacts between wild and domestic species in the Bargy massif. In this ecosystem, local populations of Alpine ibex *Capra ibex*, chamois *Rupicapra rupicapra* and domestic ruminants intermingle in Alpine pastures and have been found infected with varying prevalence between species^38^, questioning the role of these different populations in the multi-host system. Our objective was to estimate the within-species *R*_0_ as well as the *R*_0_ between all possible pairs of species in this system, to determine which hosts are involved in the maintenance and the transmission functions. This study demonstrates the value of integrating detailed empirical data on contacts within and between species into mathematical models to characterise multi-host systems at the wildlife-livestock interface.

## Results

### Relative contact rates between host species

To inform the transmission rates in our multi-host model, we used direct observations of livestock herds (cattle, sheep and goats) and wildlife individuals (ibex and chamois) in the study area during summer 2013 (Figure 1). We defined appropriate contacts for a susceptible host as all simultaneous and past (within 25 days^39^) observations of hosts of the same or different species at the same place, and derived relative contact rates between species, setting the values of each within-species contact rates to 1.

**Figure 1:**
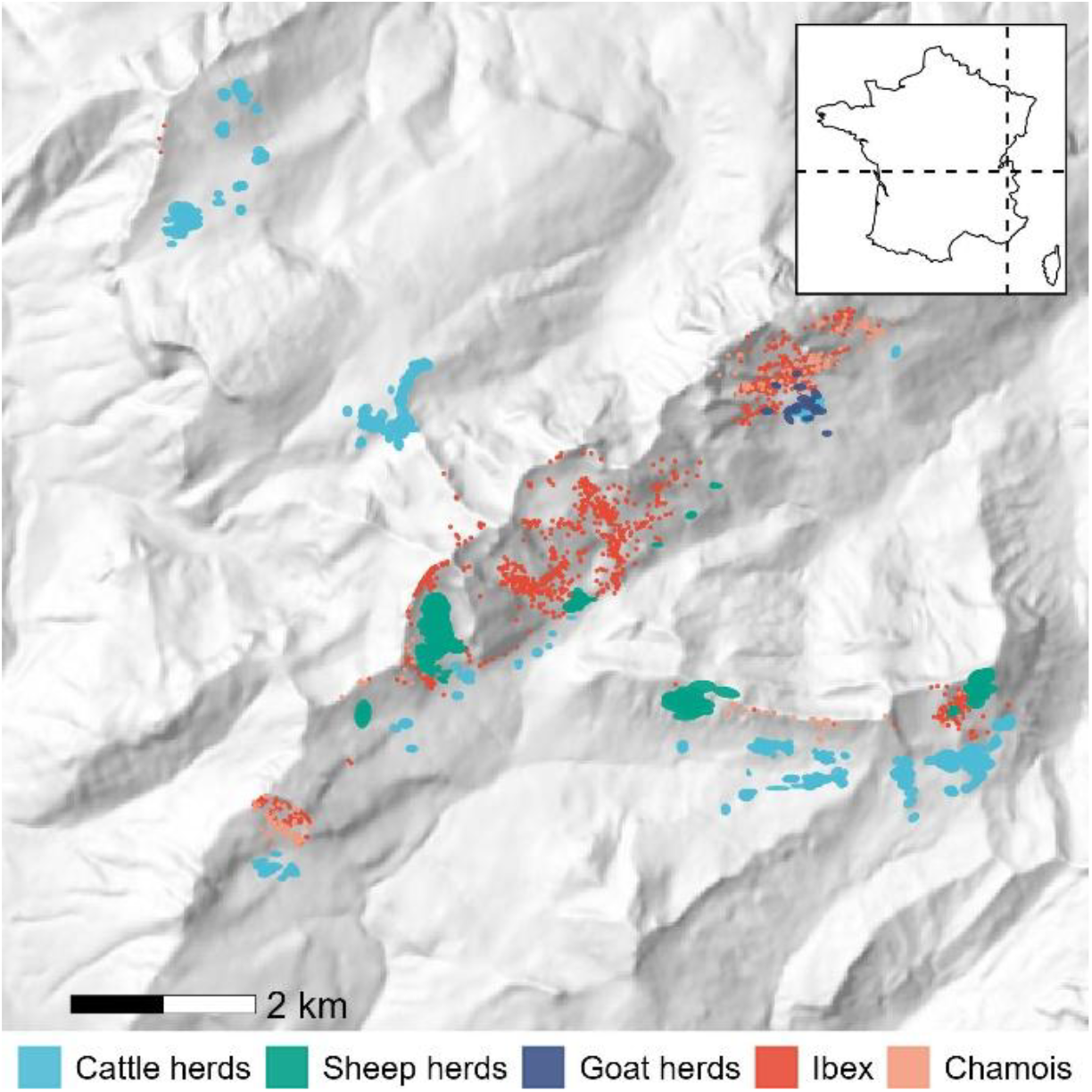
Locations of the observations of domestic ruminant herds and groups of wild individuals in 2013 in the Bargy massif, French Alps.

Table 1 and Figure S1A synthesise the estimates of the relative average contact rates. In most cases, the between-species relative contact rates were lower than one, indicating less between-species contacts than within-species contacts. There was one notable exception, where susceptible goat herds had relatively more contacts with individual ibex than with other goat herds (average relative contact rate of 3.5, 95% confidence interval - CI: [2.19-5.98]).

**Table 1:**
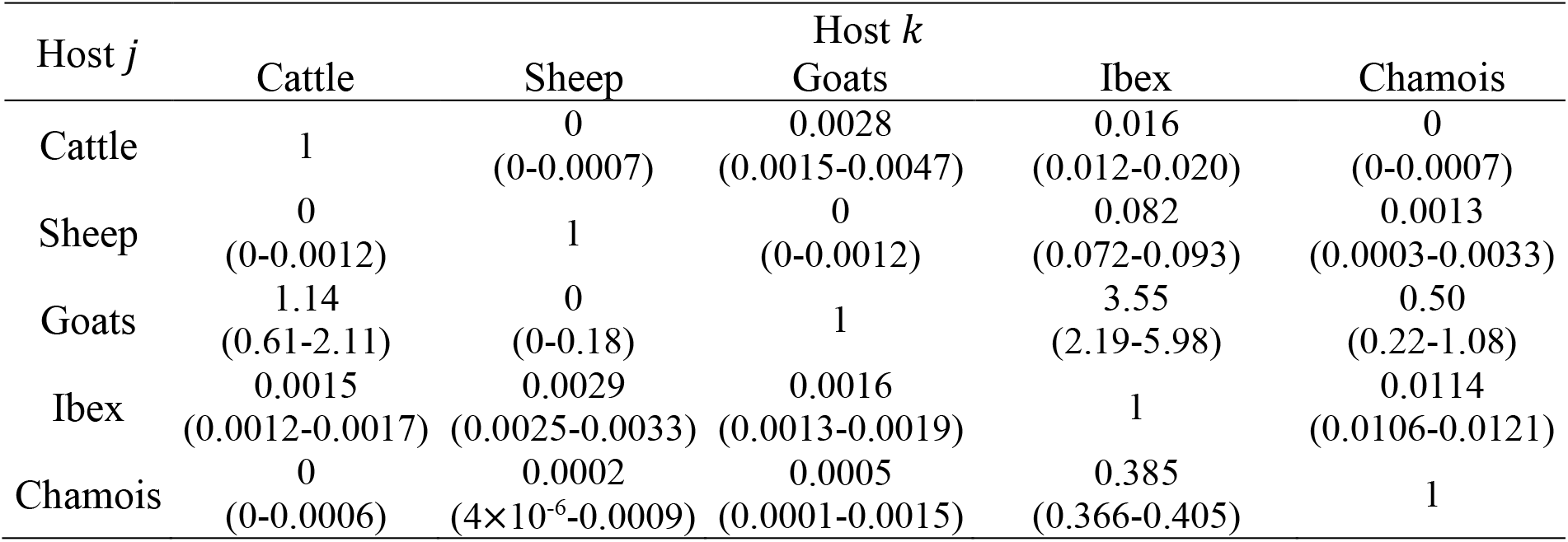
Average between-species contact rate *ω*_*jk*_ experienced by each (susceptible) wildlife individual/domestic herd of host *j* observed at a given time point relative to its average within-species contact rate *ω*_*jj*_ (set to 1). For domestic hosts (cattle, sheep and goats), the unit is a herd. For wildlife hosts (ibex and chamois), the unit is an individual animal. Results are expressed as means (95% confidence intervals).

Because infectious hosts have to precede susceptible hosts for an infectious contact to happen at a given place, the contact rates were not symmetrical. For instance, a susceptible ibex individual had very few contacts with (potentially infectious) chamois (0.0114 [0.0106-0.0121]), whereas a susceptible chamois had relatively more contacts with (potentially infectious) ibex (0.385 [0.366-0.405]). This could arise from different spatio-temporal use between the two species, where chamois use the same places as ibex but at a later time, making them more exposed to *Brucella* shed by ibex than the reverse.

### Estimates of the within-species, between-species and overall basic reproduction number

The transmission model was a multi-host version of the classical SIR (Susceptible-Infectious-Recovered) framework adapted from Nishiura *et al*. ^26^ to fit our study system. By fitting the model to prevalence data reported in previous studies^38,40^ (Figure 2), we found maximum-likelihood estimates of the average annual force of infection of 0.0062 (95% CI: 0.0004-0.0278) for cattle herds (1 positive/161 tested), 0 (0-0.044) for small ruminant herds (0 positive/45 tested), 0.093 (0.059-0.145) for ibex individuals (32 positives/80 tested), and 0.056 (0.003-0.258) for chamois individuals (1 positive/39 tested).

**Figure 2:**
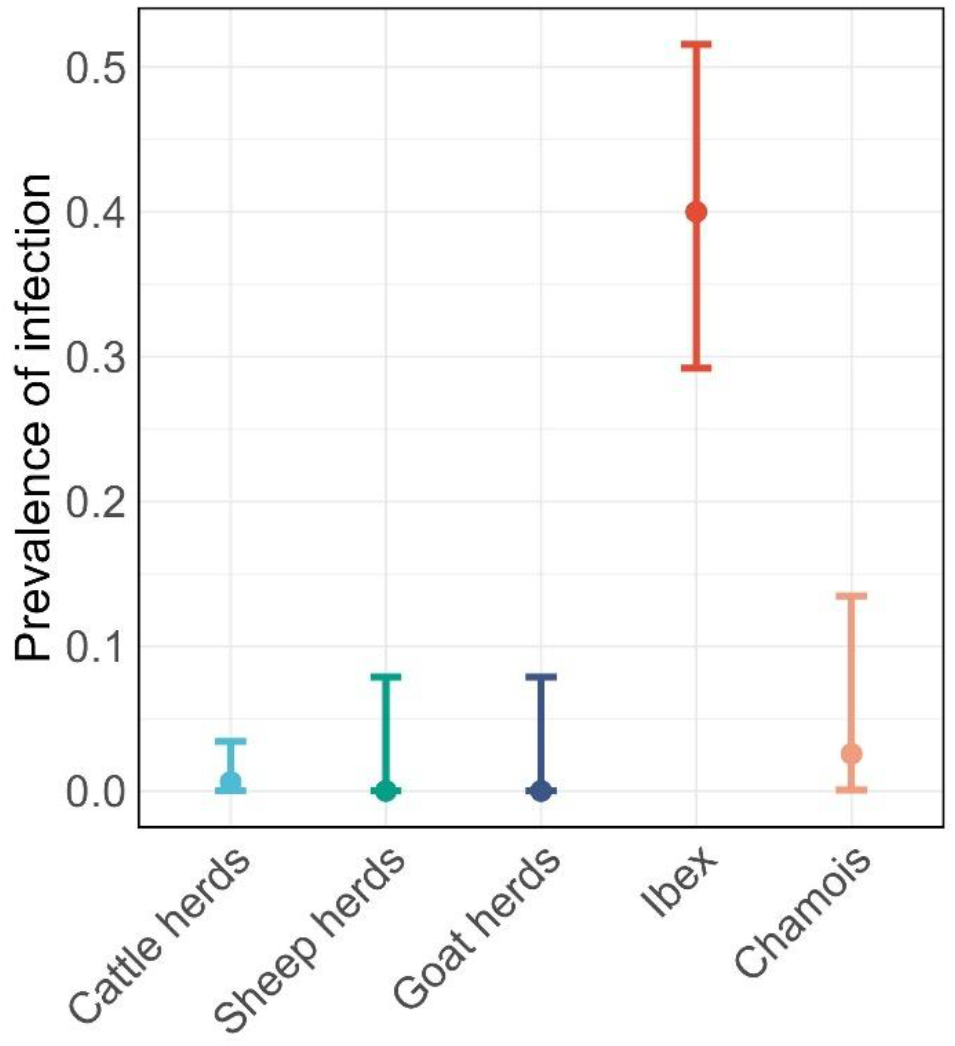
Observed prevalence of *Brucella melitensis* infection in each type of hosts in 2012-2013 in the Bargy massif, French Alps. Points indicate the mean and whiskers indicate the lower and upper bounds of the 95% confidence interval. Data from ^38,40^.

For the first time, we combined these five estimates of the force of infection with the estimates of the relative contact rates obtained from direct observations (Table 1) to calculate the intra- and interspecific transmission rates (see the WAIFW matrix in Supplementary Information and Figure S2A).

With these transmission rates, we then calculated the *R*_0_ within and between each pair of species (Table 2 and Figure S3A) as well as the overall *R*_0_. The latter was 1.67 (95% credible interval - CrI: 1.42-2.04). Ibex had the highest within-species *R*_0_, with a 95% CrI above one (1.66 [1.42-2.03]). All the other species had 95% CrI below one (Table 2 and Figure S3A). In addition, the overall *R*_0_ when excluding ibex from the system (assuming nothing else would change) had a 95% CrI below one (0.69 [0.19-0.94]).

**Table 2:** Basic reproduction number *K*_*jk*_ (average number of secondary infections resulting from one infectious host of species *k* in a totally susceptible population of species *j*) between each pair of species. For domestic hosts (cattle, sheep and goats), the unit is a herd. For wildlife hosts (ibex and chamois), the unit is an individual animal. Results are expressed as means (95% credible intervals).

| Host $j$ | Host $k$ | | | | |
| --- | --- | --- | --- | --- | --- |
|  | Cattle | Sheep | Goats | Ibex | Chamois |
| Cattle | 0.69<br>(0.19-0.94) | 0<br>(0-0.0004) | 0.0019<br>(0.0005-0.0036) | 0.0348<br>(0.0096-0.0512) | 0<br>(0-0.0002) |
| Sheep | 0<br>(0-0.0003) | 0<br>(0-0.83) | 0<br>(0-0.0003) | 0<br>(0-0.22) | 0<br>(0-0.0008) |
| Goats | 0<br>(0-0.096) | 0<br>(0-0.0029) | 0<br>(0-0.085) | 0<br>(0-0.92) | 0<br>(0-0.021) |
| Ibex | 0.0008<br>(0.0006-0.001) | 0.0015<br>(0.0012-0.0019) | 0.0008<br>(0.0007-0.0011) | 1.66<br>(1.42-2.03) | 0.0028<br>(0.0023-0.0034) |
| Chamois | 0<br>(0-0.0004) | 0.0001<br>(0-0.0007) | 0.0003<br>(0-0.0014) | 0.73<br>(0.10-1.88) | 0.28<br>(0.04-0.71) |

Regarding between-species transmission (Table 2 and Figure S3A), the *R*_0_ from ibex to chamois (0.73 [0.10-1.88]) was higher than the *R*_0_ from chamois to ibex (0.0028 [0.0023-0.0034]). Regarding cattle, the population of concern, the *R*_0_ from ibex to cattle was (0.0348 [0.0096-0.0512]), while it was 0 (0-0.0002) from chamois to cattle, as there were no observed contacts between these two species (Table 1 and Figure S1A).

The estimates of the *R*_0_ between each pair of species and of the overall *R*_0_ were very similar when assuming different durations of the survival time of *Brucella* in the environment (15 days or 49 days instead of 25 days) or when using an SIR instead of an SI model in chamois (see Supplementary Information and Figures S1-3).

## Discussion

Although theoretical frameworks on the characterization of reservoirs of pathogens in multi-host systems have received increasing attention^12–14,16,18,20^, their application to empirical situations at the wildlife-livestock or at the animal-human interface have remained relatively scarce^17,26–28,31^. In addition, these few examples either focused only on wildlife and did not include the population of concern or interest^17,26^, essential in the definition of a reservoir^12^, and/or focused only on maintenance^17,26,27^, neglecting the transmission function^13^. To inform between-species transmission rates, previous studies assumed various contact structures^26^ or sampled across the range of all possible values^17,27^. While this may be appropriate to explore the maintenance function, it is less suited to the evaluation of the transmission function because of the lack of empirical data behind the between-species transmission rates. Other studies have used spatial overlap or spatial proximity^28,31^ to inform cross-species contacts and transmission, which may be more relevant. In south-west France, this approach revealed a bovine tuberculosis maintenance community where both badger groups and cattle herds are required to maintain the pathogen, with more badger-to-cattle transmission than the opposite in recent years^31^. However, multi-host models may benefit from more direct measures of contact rates to better estimate transmission between species, particularly at the wildlife-livestock interface. Here, we integrated for the first time results from direct observations in a focal study^39,41^ into a multi-host model, building on previous works^14,26^.

Our results suggest that the basic reproduction number for *B. melitensis* transmission in the ibex population in the study area was above one (Table 2), meeting the definition of a maintenance host^12^, i.e., a population within which a pathogen can persist indefinitely even in the absence of transmission from other hosts. As the overall *R*_0_ was below 1 in the absence of ibex, they were also essential hosts^15^ in the Bargy ecosystem. These results are consistent with the hypothesis that infection was sustained in this population since at least the late 1990s, and with the observation of numerous cases of *B. melitensis* infection in the ibex population since 2012, including in recent years. This suggests that the infection still persists in this population despite a substantial decrease in seroprevalence in ibex owing to disease management interventions^42^. In contrast, the chamois population could be considered as a spillover host^14^, unable to maintain the pathogen on its own (within-species *R*_0_ below one) but still being infected by the maintenance host and potentially contributing to onward transmission, albeit in a limited way. This is in accordance with the sporadic cases observed from 2012 to 2025 in the chamois population, with one case in a found-dead animal and only four cases among >900 tested at hunting.

Because of the detection of a single infected cattle herd in 2012, our results suggest that maintenance was not possible (*R*_0_<1) in domestic ruminants in the Bargy massif under current surveillance and control measures. This result needs to be interpreted with caution, as domestic ruminants were able to maintain *Brucella* infection in the absence of transmission from other species before eradication programs were put in place and are still able to do so in areas where brucellosis is endemic^33^. In the study area, besides the first cattle outbreak, there was one other infected cattle herd detected in 2021, while no goat or sheep herds have been found infected so far. This highlights the importance of maintaining strict surveillance and control measures in domestic ruminants.

Regarding transmission from wildlife to the population of concern, i.e., cattle in the study area, our results suggest that it was possible from ibex (Tables 1 and 2), without the need of any bridge hosts^13^. In contrast, the available data did not support transmission from chamois to cattle (Tables 1 and 2), as no direct nor indirect contacts were observed between these two populations during the study period. It is possible that contacts were missed during the field observations, which are not exhaustive, as such contacts have been reported in other Alpine areas^39^. Further studies using field observations or other methods, such as global positioning system (GPS) radiocollars – already used since 2012 but only in ibex^43^, could be useful to confirm these results in the study area.

Overall, we were able to conclude that the ibex population was the essential maintenance host while also contributing to the transmission function, highlighting its predominant role in the Bargy massif ecosystem (Figure 3). Our conclusions were very robust to alternative assumptions on the temporal scale of indirect contacts (i.e., varying durations of *Brucella* survival in the environment) and on the pathogenesis of the disease in chamois.

**Figure 3:**
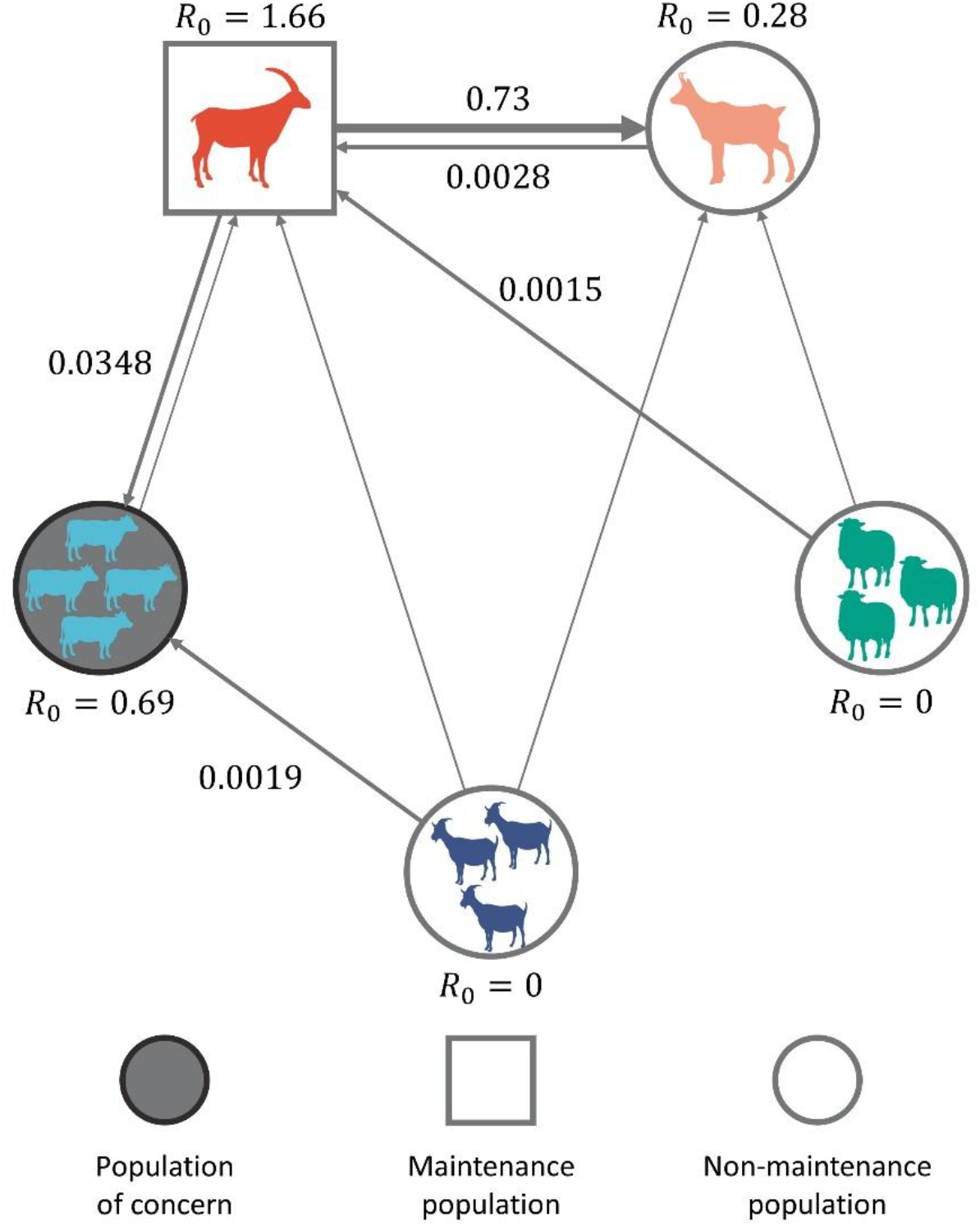
Transmission and maintenance of *Brucella melitensis* between species in the Bargy massif, French Alps. Values indicates point estimates of the basic reproduction numbers within- and between-species (Table 2). For transmission between species, only values above 10^-3^ are indicated. The size of the arrows indicates the order of magnitude of the between-species reproduction numbers. For domestic hosts (cattle, sheep and goats), the unit is a herd, represented with multiple silhouettes. For wildlife hosts (ibex and chamois), the unit is an individual animal, represented with a single silhouette. Silhouettes were obtained from phylopic.org and are under the https://creativecommons.org/publicdomain/zero/1.0 (public domain). The graphical representation of populations of concern, maintenance populations and non-maintenance populations were adapted from Haydon *et al*. ^12^ and Viana *et al*. ^18^.

These results have several implications for disease surveillance and management in the local wildlife populations with the aim to protect livestock and, ultimately, human health. In chamois, disease management interventions are probably not necessary but strict surveillance measures should continue to be implemented. In contrast, management strategies in ibex such as the ones already in place^42^ should allow for elimination of transmission in the long-term by decreasing the *R*_0_ in ibex and in the overall system below one. However, our results do not provide information on the time frame required to reach this objective. In addition, our results also support implementation of blocking tactics^12^, i.e., measures aiming at reducing contacts and transmission between wildlife and cattle to mitigate the risk of spillover. In the study area, examples of such measures include using protection dogs, keeping domestic ruminants in fences, or removing attraction points such as salt blocks. Although the risk of spillover to cattle cannot be completely eliminated as long as the bacteria is maintained in the ibex population and its environment, these measures should be encouraged to further reduce the *R*_0_ from ibex to cattle.

Beyond brucellosis in the Bargy massif, our work provides a generic and flexible approach to the study of multi-host systems when detailed empirical data on contacts within and between species are available or could be collected. Our modelling approach could for instance be applied to *B. abortus* in the Greater Yellowstone ecosystem in the USA, where genomics data strongly suggest that wild elk (*Cervus canadensis*) populations are currently maintaining the pathogen and transmitting it to livestock while wild populations of bison (*Bison bison*) are more likely spillover hosts^36^. It could also be used for *B. abortus* in several areas in south-eastern Africa, to determine in particular the roles of African buffalo (*Syncerus caffer*) and Kafue lechwe antelope (*Kobus leche kafuensis*) populations in the maintenance and the transmission functions^44^. Our framework can also be adapted to multi-host pathogens other than *Brucella* spp., such as the agent of bovine tuberculosis, another major disease at the wildlife-livestock interface where wildlife populations can play different roles according to the ecosystem^17,45^.

To be applied to other multi-host systems, the mechanistic model may have to be adapted to reflect the contact patterns between hosts and the pathogen characteristics in the ecosystem at hand. This should be relatively straightforward given the flexibility provided by SIR-like mechanistic models. For instance, we assumed frequency-dependent contact rates^26^, i.e., the number of contacts was independent of the size of the population to which the infectious host belongs. Other formulations where contact rates depend on the population size could be used^30,46^, especially when population sizes are not constant. This would require estimates of host population sizes, and ideally analysing empirical data to see whether contact rates depend on population size. As an example, direct contact rates between elks in the *B. abortus* reservoir in the Greater Yellowstone Ecosystem increased with group size but saturated in the largest groups, i.e., an intermediate between frequency- and density-dependent contact rates^46^. Contact rates could also vary seasonally, as observed for instance between GPS-equipped wild boar and domestic pig farms^47^. Time-dependent functions that can be easily added to mechanistic models can adequately describe these seasonal variations, as exemplified for Chronic Wasting Disease and contacts between white-tailed deer (*Odocoileus virginianus*) in the USA^48^. Other adaptations of the mechanistic model could include host-specific differences in infectiousness, as done for bovine tuberculosis in red deer and wild boar in Spain based on pathogen excretion data^17^, or explicitly modelling the decay rate of pathogens in the environment, as done for the same disease in badgers and cattle in Ireland^45^. One limitation to the adaptation of mechanistic models is the requirement of accurate parameter values, which will depend on the amount of available information and data in the study system. Alternatively, Fenton *et al*.^14^ developed another analytical framework relying on a limited number of parameters specifically for cases where detailed information is lacking. Our approach based on empirical contact data to estimate relative contact and transmission rates could also be integrated into their framework.

Our study relied on direct observations of wildlife and livestock (Figure 1) to infer relative contact rates (Table 1). Our approach could be similarly used for data generated through increasingly popular technologies such as GPS radiocollars, proximity loggers or camera traps^41^, making it useful in a wide range of contexts. Although contacts between wildlife and livestock are difficult to observe, these methods are facilitating their monitoring and as such have received increased attention as highlighted in a recent review^49^. Each method has its own pros and cons, summarised in Triguero-Ocaña *et al*. ^41^. Briefly, direct observation studies such as the one conducted here make it possible to record detailed information such as the number of animals, but are missing nocturnal activities and are usually restricted to small study areas and short time periods^39,41^. Camera traps can also record direct and indirect contacts between animals, but contact detection is limited to the area in front of the cameras, making them useful for monitoring aggregation points but not for estimating contact rates over large areas^41,50^. In contrast, GPS collars have high spatiotemporal resolutions, but depending on the study area external factors may lead to missed or imprecise location recordings^41^ and frequent records may be needed for epidemiological purposes^51^. Finally, proximity loggers equipped on animals can quantify direct contacts and their durations, but they lack geospatial information making them less useful for indirect contacts beyond some specific sites where fixed proximity loggers can be installed^41,52^. Examples where such methods have been used include estimating contacts between badgers and cattle with proximity loggers in the UK^52^, between cattle, red deer and wild boar using GPS in Spain^51^, and between pigs and wild boar using camera traps in Sardinia^53^. The results of such studies can be integrated into our framework to calculate relative contact rates in various systems. In addition, the results of different approaches could be compared and/or combined to evaluate the impact on the *R*_0_ estimates, which will provide guidance on which method to use for the purpose of understanding the roles of different hosts in a reservoir. Interestingly, a recent study used camera trapping to study the contacts of wildlife and humans with an Egyptian fruit bat (*Rousettus aegyptiacus*) colony infected with the zoonotic Marburg virus^54^. Results from such studies could also be used in our framework, expanding its utility not only to reservoirs at the wildlife-livestock interface but also at the animal-human interface.

The main limitation of the approach presented in this study is the requirement of both prevalence/seroprevalence data in different hosts and empirical data on cross-species contacts. To further our understanding of multi-host systems, known or suspected reservoirs of a pathogen should be studied within an integrated interface monitoring approach, combining health monitoring in the different compartments of the reservoir with data collection on within and between species contacts. Ideally, health monitoring in wildlife and livestock and/or humans should include the same components as the integrated wildlife monitoring approach, i.e., passive surveillance for the early detection of spillover events or species jumps, and active surveillance and population monitoring to obtain estimates of prevalence and population sizes and their dynamics^55^.

In conclusion, we expanded previous models used to identify the role of different species in multi-host systems. Our innovation was the explicit inclusion of empirical data on within- and between-species contacts at the wildlife-livestock interface. Our results informed the management of brucellosis, a zoonotic disease, thereby protecting both animal and human health. This new methodological approach can also be adapted to other multi-host pathogens, which will contribute to improve our understanding of these complex systems.

## Methods

### Multi-host system

The study area is located in the Bargy massif (46° N, 6.5° E, northern French Alps, Figure 1). Every year from spring to autumn, domestic cattle, sheep and goats graze in the mountain pastures, where contacts with wild ungulates can occur. In 2012, the local population of wild Alpine ibex, a protected species, was found infected by *Brucella melitensis*, after the discovery of one human case and one outbreak in a dairy cattle farm^40^. The seroprevalence in ibex captured in autumn 2012 and spring 2013 was unexpectedly high^38^ at 40% (95% CI: 29.2%-51.6%, *n*=80, Figure 2). The close genetic proximity between isolated strains in ibex and cattle in 2012 and the one isolated in 1999 from the last local bovine case led to suspect the persistence of *B. melitensis* in ibex since the late 1990s, and its subsequent spillover to cattle in 2012^37,38^.

Beyond the first infected cattle farm, no other herd was found infected among the remaining 160 cattle herds and 45 small ruminant herds that grazed in the Bargy massif during the summer of 2012^38^. Surveillance among hunted wild ruminants during winter 2012-2013 in the massif revealed one positive chamois (*n*=39) in both serology and bacteriology, but no infected red deer or roe deer (*Capreolus capreolus*)^38^. In addition, red deer and roe deer do not use the same areas as the other species, preferring lower altitudes.

Therefore, in our model, we included two wild species: ibex and chamois, and three domestic species: cattle, sheep and goats, for which both prevalence and observation data were available. Humans were not included given their very different contamination route in the study area, i.e., through the consumption of unpasteurized dairy products^37,40^.

### Estimates of the relative contact rates

To estimate the relative contact rates at the wildlife-livestock interface, we used direct observations of wildlife and livestock groups^39^. Field observations of nine sites chosen for their representativity and their accessibility were carried out between 19 June and 6 August 2013^56^. For each site, the observations lasted 1-3 consecutive days depending on the weather and were repeated 1-3 times during the study period. During each observation session, pairs of observers were stationed at a viewpoint from which they could observe the mountain pastures using binoculars and spyglasses. Every 30 minutes, from 6am to 10am and from 5pm to 9pm, and every two hours in between, observers reported the positions of wild and domestic ruminants on a map. Individual or groups of wild ruminants were represented by points at the centroid of the group, while domestic herds were represented by polygons, encompassing all the animals belonging to the herd. For each point or polygon, the date, time, species and number of individuals (for wildlife) was recorded. Points and polygons were manually digitised into shapefiles using the GIS software MapInfo. For wildlife, a buffer of 20 m around the point was added to account for the precision of recorded locations^56^. There was a total of 874 observations of domestic ruminant herds (cattle: 610, sheep: 239, goats: 25), 1,302 observations of ibex groups (4,596 individuals) and 225 observations of chamois groups (517 individuals – Figure 1).

The main transmission route of *B. melitensis* in domestic and wild ruminants is by direct or indirect contact with infectious materials shed during abortion or birth^57,58^. Therefore, appropriate contacts for *Brucella* transmission to be possible between one susceptible and one infectious host include both direct contacts between animals at the same time and place, and contacts with contaminated environment. We defined a contact as two observations that were less than 25 days apart (critical time *sensu* Bacigalupo *et al*.^49^), based on the estimated survival time of *Brucella* in mountain pastures^39^. Alternative values (15 days and 49 days) were also evaluated (see *Sensitivity analysis*). We also assumed that a contact occurred when herd polygons and/or wildlife buffer intersected (critical space *sensu* Bacigalupo *et al*.^49^). We defined the contact rate *c*_*jk*_, i.e., the average number of wildlife individuals/domestic herds of species *k* contacted by each (susceptible) wildlife individual/domestic herd of species *j* observed at a given time point, as:

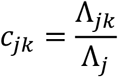

where *N*_*j*_∼*Poisson*(Λ_*j*_) is the total number of observations of wildlife individuals/domestic herds of species *j*, and (∑ *n*_*jk*_)∼*Poisson*(Λ_*jk*_) is the total number of observations of wildlife individuals/domestic herds of species *j* in contact with wildlife individuals/domestic herds of species *k*. In our context, a contact meant that the location of species *k* intersected with the location of species *j* within the 25 days that preceded the observation of species *j* at that location. We pooled all observations, averaging the contact rates over the whole area and study period.

The observations were restricted temporally to the summer and at daytime when the weather allowed good visibility, and spatially to nine open areas with good viewpoints. To limit the risk of potential biases, we set the values of each within-species contact rate to 1 and derived relative contact rates rather than absolute values, assuming that the relative values of the contact rates would remain the same if it was possible to observe all contacts. We calculated the relative contact rate *ω*_*jk*_ as:

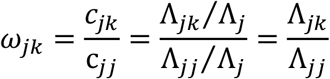

The mean and 95% confidence interval of *ω*_*jk*_ (Table 1) were calculated with the function “rateratio.test” from the homonym package^59^ in the statistical software R 4.3.3^60^.

### Transmission model

We developed a multi-host model of *B. melitensis* transmission with *n* = 5 different host species: cattle, sheep, goats, ibex and chamois. To match both prevalence data and intra- and interspecific contact data, the epidemiological unit was at the individual level for ibex and chamois while it was at the herd level for domestic ruminants.

For domestic ruminant herds, we used an SIS (susceptible-infectious-susceptible) model, considering that infected herds should be depopulated immediately after detection. The following set of differential equations was used:

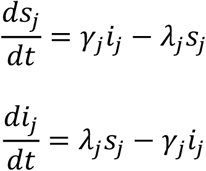

where *j* is the host species (cattle, sheep or goats), *λ*_*j*_ is the force of infection (i.e., the average annual rate at which susceptible herds become infected, regardless of the source of infection), and 1/*γ*_*j*_ is the average duration of the infectious period (i.e., between onset of infection and depopulation). We assumed that herds were quickly repopulated after depopulation and appropriate cleaning and disinfection. We used proportions of susceptible (*s*_*j*_) and infectious (*i*_*j*_) herds (hence the use of lowercase instead of capital letters), assuming *s*_*j*_ + *i*_*j*_ = 1 at any time *t*. In the 2012 infected cattle farm, detection and whole-herd depopulation most likely occurred one year after the herd was infected^54^. We therefore assumed an approximate one-year delay (1/*γ*_*j*_) between onset of infection and herd depopulation. In the absence of outbreak in small ruminants, we assumed a similar value for sheep and goat herds.

For ibex individuals, we used an SIR (susceptible-infectious-recovered) model^58^, neglecting the latent period which is usually short (about three weeks) in ruminants^61^. Despite the presence of gross lesions (arthritis, orchitis) in some animals, the available evidence does not currently support brucellosis-related mortality in the ibex population^58,62^. Therefore, the following set of differential equations was used:

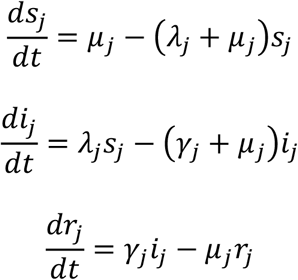

where *j* is the host species (here Alpine ibex), *λ*_*j*_ is the force of infection (i.e., the average annual rate at which susceptible individuals become infected, regardless of the source of infection), 1/*γ*_*j*_ is the average duration of the infectious period, and *µ*_*j*_ is the annual natural mortality rate. We used proportions of susceptible (*s*_*j*_), infectious (*i*_*j*_) and recovered (*r*_*j*_) individuals, assuming *s*_*j*_ + *i*_*j*_ + *r*_*j*_ = 1 at any time *t*. We thus assumed that the annual birth rate was identical to the natural mortality rate *µ*_*j*_, which was estimated at 0.13 per year (C. Toïgo, pers. comm.). The recovery rate *γ*_*j*_ was previously estimated at 0.17 per year^58^.

Finally, we used a SI (susceptible-infectious) model for chamois individuals, considering the high pathogenicity of brucellosis in this species^63–66^, which generally leads to death in a few months^66^. Therefore, the following set of differential equations was used:

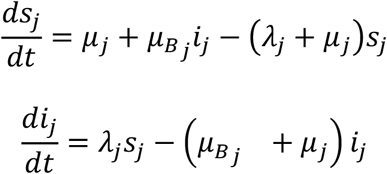

where *j* is the host species (here chamois), *λ*_*j*_ is the force of infection (i.e., the average annual rate at which susceptible individuals become infected, regardless of the source of infection), *µ*_*Bj*_ is the brucellosis-related mortality, and *µ*_*j*_ is the annual natural mortality rate. We used proportions of susceptible (*s*_*j*_) and infectious (*i*_*j*_) individuals, assuming *s*_*j*_ + *i*_*j*_ = 1 at any time *t*. We thus assumed that the annual birth rate was identical to the natural mortality rate *µ*_*j*_ and that all infected individuals died and were replaced by births of susceptible individuals. The natural mortality rate *µ*_*j*_ was estimated at 0.14 per year (C. Toïgo, pers. comm.), and we assumed a disease-induced mortality *µ*_*Bj*_ of 2 per year, corresponding to an average duration of the infectious period of 6 months. As an alternative, we assessed how an SIR model (similar to ibex) in chamois would impact our results (see *Sensitivity analysis*).

### Estimates of the force of infection

We focused on the situation up to the summer of 2013, before intense management operations started to be implemented in ibex. In particular, a drastic culling operation involving 251 individuals was conducted in autumn 2013-spring 2014^43^. We assumed that demographic and infection dynamics were on average at equilibrium up to the summer of 2013^58,67^, which was supported by the reintroduction of the ibex population in the 1970s and demographic data suggesting limited growth in 2013^67^, and by the presence of *B. melitensis* in ibex for more than a decade^37^. We also averaged contacts and transmission over a yearly time step, consistent with the relatively slow-spreading nature of *Brucella* and the lifespan of the hosts, which allowed us to overlook potential seasonal oscillations for simplicity.

As in Nishiura *et al*. ^26^, we estimated the average annual force of infection *λ*_*j*0_ of host *j* in the endemic steady state by minimising the negative log-likelihood for each host species, using the “optimize” function in R and assuming:

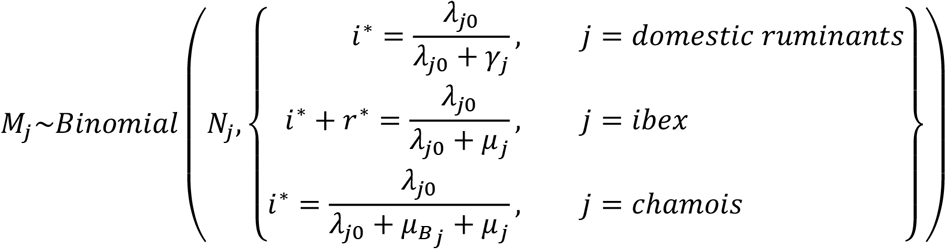

where *M*_*j*_ was the number of infected out of *N*_*j*_ tested herds/individuals. The 95% profile likelihood confidence intervals (CIs) were determined by including all the values of the force of infection that produced negative log-likelihoods below 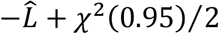, where 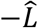 was the overall minimum negative log-likelihood^68^. Because the prevalence data (Figure 2) was pooled for small ruminant herds (no distinction between goat herds and sheep herds)^38^, we obtained the same force of infection for both sheep and goats.

### Estimates of the transmission rates and the basic reproduction number

Assuming frequency-dependent transmission, the force of infection at equilibrium *λ*_*j*0_ was^26^:

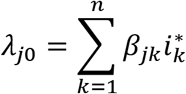

where *β*_*jk*_ was the transmission rate, i.e., the average number of individuals/herds of species *k* effectively contacted by each (susceptible) individual/herd of species *j* per year. With the five estimates of the force of infection, it would be possible to calculate five transmission rates. However, there are as many transmission rates as there are possible pairs of species in the ‘who acquires infection from whom’ (WAIFW) matrix^15^, i.e., *n* × *n* transmission rates where *n* is the number of hosts (*n* = 5 in our case). Therefore, it was necessary to reduce the number of parameters to estimate to no more than *n* values.

The transmission rate *β*_*jk*_ is the product of two components, i.e. the contact rate 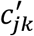 (average number of individuals/herds of species *k* contacted by each individual/herd of species *j* per year) and the probability *ν*_*jk*_ of successful transmission given contact between an infectious and a susceptible host^30^. Here, we did not have access to the average number of intra- and interspecific contacts per year 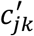, which would require to continuously follow every host, but only to the average number of intra- and interspecific contacts per observation *c*_*jk*_. Therefore, we assumed that their relative values were maintained, e.g., if the observation data shows that an ibex has 10 times more contacts with cattle herds than with sheep herds, then the same proportion will be true for the average number of contacts per year. We further assumed for parameter identifiability that the probability of successful transmission given contact was the same for all infectious hosts *k* on a given susceptible host *j, i*. *e*., *ν*_*jk*_ = *ν*_*jj*_ for all *k*. Thus, we assumed that host-specific differences in bacterial shedding was negligible, and that the probability of transmission relied only on host-specific differences in susceptibility.

This allowed us to assume that 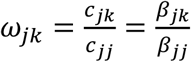, where *ω*_*jk*_ was the relative contact rate calculated from the observations (Table 1). Therefore, the force of infection was re-written as:

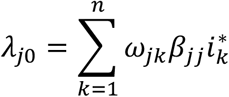

Resolving this equation, it was possible to calculate the intra- and interspecific transmission rates:

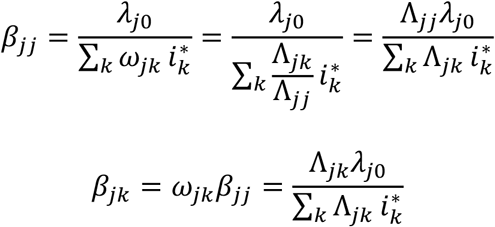

where *λ*_*j*0_, Λ_*jk*_ and 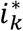 were all calculated at previous steps.

This allowed us to obtain each element *β*_*jk*_ of the WAIFW matrix, noted *T*. The derivation of the corresponding confidence intervals was not straightforward as the transmission rates *β* combine several parameter estimates. Therefore, we simulated 100 000 values of 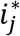 (or 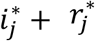 for ibex) and Λ_*jk*_, assuming their distribution was *Beta*(*M*_*j*_ + 0.5, *N*_*j*_ − *M*_*j*_ + 0.5) and *Gamma* ((∑ *n*_*jk*_) + 0.5, 1), respectively, which is equivalent to using the Jeffreys interval^69,70^ for a binomial proportion 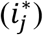 or a Poisson rate(Λ_*jk*_). For each simulation, we calculated *λ*_*j*0_, Λ_*jk*_ and 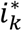 and the corresponding *β*_*jk*_. The credible interval (CrI) was then calculated as the equal-tailed 95% interval of the distribution of the 100 000 values of *β*_*jk*_.

Finally, the next-generation matrix *K* is given by *K* = −*T*Σ^−1^, where Σ is the transition matrix, i.e. the *n* × *n* matrix corresponding to recovery and death of state *i*^24^. Here, we obtain:

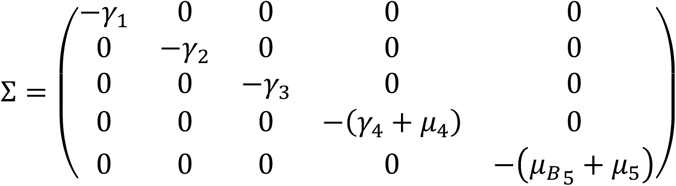

Therefore, −Σ^−1^ describes the average duration that an individual/herd will stay in the infectious state *i*^24^. The next-generation matrix was:

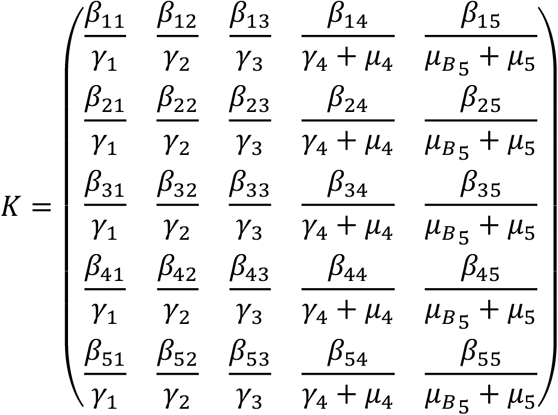

Each element of *K* can be interpreted in a similar way as the basic reproduction number^25^: *K*_*jk*_ is the expected number of secondary cases in host *j* that would arise from the introduction of one infectious host *k* in a susceptible population of host *j*. The overall basic reproduction number *R*_0_ in the multi-host system is the dominant eigenvalue of the next-generation matrix^24^.

Finally, we also calculated the host-specific reproduction number *U*_*m*_ (measuring the expected number of secondary infections in hosts *m* resulting from an average infectious individual of hosts *m* in the absence of other hosts) and the host-excluded reproduction number *Q*_*m*_ (measuring the expected number of secondary infections in hosts other than *m* resulting from an average infectious individual of hosts other than *m* in the absence of hosts *m*)^25,26^, as follows:

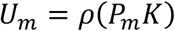

and

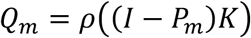

where 1 ≤ *m* ≤ *n, I* is the *n* × *n* identity matrix, *P* is the *n* × *n* projection matrix defined by *P*_*jj*_ = 1 when *j* ∈ *m, P*_*jj*_ = 0 when *j* ∉ *m* and *P*_*jk*_ = 0 when *j* ≠ *k*.

### Sensitivity analysis

We assessed the sensitivity of our results to two critical model assumptions: the survival time of *Brucella* in pastures, used to estimate the relative contact rates, and the disease-induced mortality in chamois. For the survival time of *Brucella* in the mountain environment, we evaluated two values in addition to the baseline value of 25 days: a lower value of 15 days^39^ and a higher value taking into account all possible indirect contacts during the observation period (maximum of 49 days). The assumption that all infectious chamois died after a few month was based on past observations of a few sporadic cases of *B. abortus* and *B. melitensis* in chamois in Europe, which displayed very severe lesions suggesting disease-induced mortality^63–66^. High disease-induced mortality in chamois could also explain why self-sustained persistence of *Brucella* in the absence of transmission from domestic species have never been observed in this species^64,67^. However, some individual chamois do not seem to display symptoms^64^. In the study area, there were only a few cases, including a single case in found-dead animals. Therefore, the information was scarce in chamois compared to the available knowledge in ibex, which led us to evaluate an alternative SIR model for chamois.

In the absence of specific data, we used the same recovery rate in chamois than in ibex (0.17 per year^58^). The reality is probably in-between these two model formulations, but further data would be required to gain a better understanding of recovery and brucellosis-induced mortality in chamois.

## Acknowledgments

The authors are grateful to all the people involved in collecting field data and running laboratory analyses. The authors wish to thank the “Direction Départementale des Territoires” of Haute-Savoie, France, for providing the data on direct observations of wildlife and livestock groups. This study was funded by the Veterinary School of Toulouse (project “INTEREPI”, BQR 2024).

## Author contributions

SL designed the study and acquired funding to support this project. SL supervised CM, and both developed the model and performed the analyses, with input from all authors. PB, DG, and EP participated to data collection and curation. All authors contributed to the interpretation of the results. SL drafted the manuscript and all authors substantially revised it. All authors approved the final version of the manuscript.

## Competing interests

The authors declare no competing interests.

## Notes

### Competing Interest Statement

The authors have declared no competing interest.

## References

1. Taylor, L. H., Latham, S. M. & Woolhouse, M. E. J. Risk factors for human disease emergence. Philos. Trans. R. Soc. Lond., B, Biol. Sci. 356, 983–989 (2001).

2. Woolhouse, M. E. J. & Gowtage-Sequeria, S. Host range and emerging and reemerging pathogens. Emerg. Infect. Dis. 11, 1842–1847 (2005).

3. Cleaveland, S., Laurenson, M. K. & Taylor, L. H. Diseases of humans and their domestic mammals: pathogen characteristics, host range and the risk of emergence. Philos. Trans. R. Soc. Lond., B, Biol. Sci. 356, 991–999 (2001).

4. Leroy, E. M. et al. Fruit bats as reservoirs of Ebola virus. Nature 438, 575–576 (2005).

5. Hayes, B., Vergne, T., Rose, N., Mortasivu, C. & Andraud, M. A multi-host mechanistic model of African swine fever emergence and control in Romania. Nat. Commun. 17, 2659 (2026).

6. Miller, B. J. Why unprecedented bird flu outbreaks sweeping the world are concerning scientists. Nature 606, 18–19 (2022).

7. Jones, K. E. et al. Global trends in emerging infectious diseases. Nature 451, 990–993 (2008).

8. Daszak, P., Cunningham, A. & Hyatt, A. D. Emerging infectious diseases of wildlife-Threats to biodiversity and human health. Science 287, 443–449 (2000).

9. Wiethoelter, A. K., Beltrán-Alcrudo, D., Kock, R. & Mor, S. M. Global trends in infectious diseases at the wildlife–livestock interface. Proc. Natl. Acad. Sci. U.S.A. 112, 9662–9667 (2015).

10. Karmacharya, D., Herrero-García, G., Luitel, B., Rajbhandari, R. & Balseiro, A. Shared infections at the wildlife–livestock interface and their impact on public health, economy, and biodiversity. Anim. Front. 14, 20–29 (2024).

11. Woolhouse, M. E. J., Taylor, L. H. & Haydon, D. T. Population biology of multihost pathogens. Science 292, 1109–1112 (2001).

12. Haydon, D. T., Cleaveland, S., Taylor, L. H. & Laurenson, M. K. Identifying reservoirs of infection: a conceptual and practical challenge. Emerg. Infect. Dis. 8, 1468–1473 (2002).

13. Caron, A., Cappelle, J., Cumming, G. S., de Garine-Wichatitsky, M. & Gaidet, N. Bridge hosts, a missing link for disease ecology in multi-host systems. Vet. Res. 46, 1–11 (2015).

14. Fenton, A., Streicker, D. G., Petchey, O. L. & Pedersen, A. B. Are all hosts created equal? Partitioning host species contributions to parasite persistence in multihost communities. Am. Nat. 186, 610–622 (2015).

15. Webster, J. P., Borlase, A. & Rudge, J. W. Who acquires infection from whom and how? Disentangling multi-host and multi-mode transmission dynamics in the ‘elimination’ era. Philos. Trans. R. Soc. Lond., B, Biol. Sci. 372, 20160091 (2017).

16. Fenton, A. & Pedersen, A. B. Community epidemiology framework for classifying disease threats. Emerg. Infect. Dis. 11, 1815–1821 (2005).

17. Santos, N. et al. Quantification of the animal tuberculosis multi-host community offers insights for control. Pathogens 9, 421 (2020).

18. Viana, M. et al. Assembling evidence for identifying reservoirs of infection. Trends Ecol. Evol. 29, 270–279 (2014).

19. Wilber, M. Q., DeMarchi, J., Fefferman, N. H. & Silk, M. J. High prevalence does not necessarily equal maintenance species: Avoiding biased claims of disease reservoirs when using surveillance data. J. Anim. Ecol. 91, 1740–1754 (2022).

20. Hallmaier-Wacker, L. K., Munster, V. J. & Knauf, S. Disease reservoirs: from conceptual frameworks to applicable criteria. Emerg. Microbes Infect. 6, e79 (2017).

21. Lembo, T. et al. Exploring reservoir dynamics: a case study of rabies in the Serengeti ecosystem. J. Appl. Ecol. 45, 1246–1257 (2008).

22. Léger, E. et al. Prevalence and distribution of schistosomiasis in human, livestock, and snail populations in northern Senegal: a One Health epidemiological study of a multi-host system. Lancet Planetary Health 4, e330–e342 (2020).

23. de Wit, M. M. et al. Silent reservoir species are shaping the emergence of Usutu virus.Nat. Ecol. Evol. 1–12 (2026) doi:10.1038/s41559-025-02973-4.

24. Diekmann, O., Heesterbeek, J. A. P. & Roberts, M. G. The construction of next-generation matrices for compartmental epidemic models. J. R. Soc. Interface 7, 873–885 (2010).

25. Roberts, M. G. & Heesterbeek, J. A. P. A new method for estimating the effort required to control an infectious disease. Proc. R. Soc. B 270, 1359–1364 (2003).

26. Nishiura, H., Hoye, B., Klaassen, M., Bauer, S. & Heesterbeek, H. How to find natural reservoir hosts from endemic prevalence in a multi-host population: a case study of influenza in waterfowl. Epidemics 1, 118–128 (2009).

27. Rudge, J. W. et al. Identifying host species driving transmission of schistosomiasis japonica, a multihost parasite system, in China. Proc. Natl. Acad. Sci. U.S.A. 110, 11457–11462 (2013).

28. Funk, S., Nishiura, H., Heesterbeek, H., Edmunds, W. J. & Checchi, F. Identifying transmission cycles at the human-animal interface: the role of animal reservoirs in maintaining gambiense Human African Trypanosomiasis. PLoS Comput. Biol. 9, e1002855 (2013).

29. Buhnerkempe, M. G. et al. Eight challenges in modelling disease ecology in multi-host, multi-agent systems. Epidemics 10, 26–30 (2015).

30. Begon, M. et al. A clarification of transmission terms in host-microparasite models: numbers, densities and areas. Epidemiol. Infect. 129, 147–153 (2002).

31. Bouchez-Zacria, M. et al. Analysis of a multi-type resurgence of Mycobacterium bovis in cattle and badgers in Southwest France, 2007-2019. Vet. Res. 54, 1–20 (2023).

32. Hayes, B. H., Vergne, T., Andraud, M. & Rose, N. Mathematical modeling at the livestock-wildlife interface: scoping review of drivers of disease transmission between species. Frontiers in Veterinary Science 10, (2023).

33. Godfroid, J. Brucellosis in livestock and wildlife: zoonotic diseases without pandemic potential in need of innovative one health approaches. Arch. Public Health 75, 1–6 (2017).

34. Pappas, G., Papadimitriou, P., Akritidis, N., Christou, L. & Tsianos, E. V. The new global map of human brucellosis. Lancet Infect. Dis. 6, 91–99 (2006).

35. Laine, C. G., Johnson, V. E., Scott, H. M. & Arenas-Gamboa, A. M. Global estimate of human brucellosis incidence. Emerg. Infect. Dis. 29, 1789–1797 (2023).

36. Kamath, P. L. et al. Genomics reveals historic and contemporary transmission dynamics of a bacterial disease among wildlife and livestock. Nat. Commun. 7, 11448 (2016).

37. Mick, V. et al. Brucella melitensis in France: persistence in wildlife and probable spillover from Alpine ibex to domestic animals. PLoS ONE 9, e94168 (2014).

38. Garin-Bastuji, B. et al. Reemergence of Brucella melitensis in wildlife, France. Emerg. Infect. Dis. 20, 1570–1571 (2014).

39. Richomme, C., Gauthier, D. & Fromont, E. Contact rates and exposure to inter-species disease transmission in mountain ungulates. Epidemiol. Infect. 134, 21–30 (2006).

40. Mailles, A. et al. Re-emergence of brucellosis in cattle in France and risk for human health. Euro Surveill. 17, 1–3 (2012).

41. Triguero-Ocaña, R., Vicente, J., Lavelle, M. & Acevedo, P. Collecting data to assess the interactions between livestock and wildlife. in Diseases at the Wildlife-Livestock Interface: Research and Perspectives in a Changing World (eds. Vicente, J., Vercauteren, K. C. & Gortázar, C.) 307–338 (Springer International Publishing, Cham, 2021).

42. Calenge, C. et al. Estimating disease prevalence and temporal dynamics using biased capture serological data in a wildlife reservoir: the example of brucellosis in Alpine ibex (Capra ibex). Prev. Vet. Med. 187, 1–12 (2021).

43. Marchand, P. et al. Sociospatial structure explains marked variation in brucellosis seroprevalence in an Alpine ibex population. Sci. Rep. 7, 15592 (2017).

44. Simpson, G. et al. Brucellosis in wildlife in Africa: a systematic review and meta-analysis. Sci. Rep. 11, 5960 (2021).

45. Chang, Y. et al. Inferring bovine tuberculosis transmission between cattle and badgers via the environment and risk mapping. Front. Vet. Sci. 10, 1233173 (2023).

46. Cross, P. C. et al. Female elk contacts are neither frequency nor density dependent.Ecology 94, 2076–2086 (2013).

47. Morelle, K. et al. Spatio-temporal patterns and risk factors of wild boar–pig farm contact across Europe. J. Appl. Ecol. 63, e70314 (2026).

48. Williams, D. M., Quinn, A. C. D. & Porter, W. F. Informing disease models with temporal and spatial contact structure among GPS-collared individuals in wild populations. PLOS ONE 9, e84368 (2014).

49. Bacigalupo, S. A., Dixon, L. K., Gubbins, S., Kucharski, A. J. & Drewe, J. A. Towards a unified generic framework to define and observe contacts between livestock and wildlife: a systematic review. PeerJ 8, e10221 (2020).

50. Triguero-Ocaña, R., Vicente, J., Palencia, P., Laguna, E. & Acevedo, P. Quantifying wildlife-livestock interactions and their spatio-temporal patterns: Is regular grid camera trapping a suitable approach? Ecol. Indic. 117, 106565 (2020).

51. Herraiz, C. et al. Movement-driven modelling reveals new patterns in disease transmission networks. Journal of Animal Ecology 93, 1275–1287 (2024).

52. Drewe, J. A. et al. Performance of proximity loggers in recording intra-and inter-species interactions: a laboratory and field-based validation study. PLOS ONE 7, e39068 (2012).

53. Cadenas-Fernández, E. et al. Free-ranging pig and wild boar interactions in an endemic area of African swine fever. Front. Vet. Sci. 6, 376 (2019).

54. Atukwatse, B. et al. Multi-species foraging on a Marburg virus bat reservoir. Current Biology 36, R322–R323 (2026).

55. Cardoso, B. et al. Stepping up from wildlife disease surveillance to integrated wildlife monitoring in Europe. Res. Vet. Sci. 144, 149–156 (2022).

56. Gauthier, D. et al. Evaluation du risque de contamination domestique-sauvage en alpage : exemple de la brucellose du Bouquetin du Bargy. in 32èmes rencontres du GEEFSM (Olivone, 2014).

57. Diaz-Aparicio, E. Epidemiology of brucellosis in domestic animals caused by Brucella melitensis, Brucella suis and Brucella abortus. Rev. Sci. Tech. Off. Int. Epizoot. 32, 43–51 (2013).

58. Lambert, S. et al. An individual-based model to assess the spatial and individual heterogeneity of Brucella melitensis transmission in Alpine ibex. Ecol. Modell. 425, 109009 (2020).

59. Fay, M. rateratio.test: Exact Rate Ratio Test. (2022).

60. R Core Team. R: a language and environment for statistical computing. R Foundation for Statistical Computing (2024).

61. European Commission. Brucellosis in Sheep and Goats (Brucella Melitensis). 89 https://food.ec.europa.eu/document/download/e377914a-810a-4afe-a84c-147f6f12a636_en?filename=sci-com_scah_out59_en.pdf&prefLang=fr (2001).

62. Lambert, S. et al. Combining seroprevalence and capture-mark-recapture data to estimate the force of infection of brucellosis in a managed population of Alpine ibex. Epidemics 38, 100542 (2022).

63. Bouvier, G., Burgisser, H. & Schneider, P. A. La brucellose du chamois. in Les maladies des ruminants sauvages de la Suisse 111–113 (Lausanne, Suisse, 1958).

64. Ferroglio, E., Gennero, S., Rossi, L. & Tolari, F. Monitoraggio di un focolaio di brucellosi nel camoscio alpino. Journal of Mountain Ecology 7, 229–232(2003).

65. Garin-Bastuji, B., Oudar, J., Richard, Y. & Gastellu, J. Isolation of Brucella melitensis biovar 3 from a chamois (Rupicapra rupicapra) in the southern French alps. J. Wildl. Dis. 26, 116–118 (1990).

66. Gauthier, D. Brucellosis in free ranging chamois (Rupicapra rupicapra): relationships with mountain cattle breeding. in Third Conference of the European Wildlife Disease Association (Edinburgh, Scotland, 1998).

67. ANSES. Mesures de Maîtrise de La Brucellose Chez Les Bouquetins Du Bargy. 194 https://www.anses.fr/fr/system/files/SANT2014sa0218Ra.pdf (2015).

68. Bolker, B. M. Ecological Models and Data in R. (Princeton University Press, 2008). doi:10.2307/j.ctvcm4g37.

69. Brown, L. D., Cai, T. T. & DasGupta, A. Interval estimation for a binomial proportion. Statist. Sci. 16, 101–133 (2001).

70. Laud, P. ratesci: Confidence Intervals and Tests for Comparisons of Binomial Proportions or Poisson Rates. (2025).

