## Supplementary Information for "Integrating multi-host modelling with empirical wildlife-livestock contacts reveals an essential population in a pathogen reservoir"

This document is a supplementary information to the article entitled "Using data on interspecies contacts to identify reservoirs of infection between wildlife and livestock". It was produced using R Markdown to ease reproducibility.

### 1 Main results

#### 1.1 Relative contact rates between host species

The data on direct observations of wildlife and livestock groups were provided by the "Direction Départementale des Territoires" from Haute-Savoie, France. There were two shapefiles (one for domestic ruminant herds and one for groups of wild individuals) recording the date, time, species and number of individuals (for wildlife) for each observation. Individual or groups of wild ruminants were represented by points, while domestic herds (cattle, sheep and goats) were represented by polygons, with corresponding coordinates. For wildlife, a buffer of 20 meters around the point was added to account for the precision of recorded locations.

```
# Read data for domestic ruminants
domestic <- read_sf("data/domestic.gpkg")

# Read data for wildlife
wild <- read_sf("data/wild.gpkg")

# Subset by species
ibex <- subset(wild, wild$species == "Ibex")
ibex <- ibex[!is.na(ibex$number), ] # remove when number not reported (n=2)
chamois <- subset(wild, wild$species == "Chamois")
cattle <- subset(domestic, domestic$species == "Cattle herds")
goats <- subset(domestic, domestic$species == "Goat herds")
sheep <- subset(domestic, domestic$species == "Sheep herds")
```

Appropriate contacts for *Brucella* transmission to be possible between one susceptible and one infectious host include both direct contacts between animals at the same time and place, and contacts with contaminated environment. Therefore, in our context, a contact meant that the location of species $k$ intersected with the location of (susceptible) species $j$ within the 25 days that preceded the observation of species $j$ at that location. We calculated the relative contact rate $\omega_{jk}$ as the total number of observations of wildlife individuals/domestic herds of species $j$ in contact with wildlife individuals/domestic herds of a different species $k$ ($\sum n_{jk}) \sim Poisson(\Lambda_{jk})$) divided by the total number of observations of wildlife individuals/domestic herds of species $j$ in contact with wildlife individuals/domestic herds of the same species $j$ ($\sum n_{jj}) \sim Poisson(\Lambda_{jj})$). The 95% confidence interval (CI) of $\omega_{jk}$ was calculated with the function `rateratio.test`.

```
# Parameters
time_limit <- 25 # (days)
species <- c("cattle", "sheep", "goats", "ibex", "chamois")

# Initialisation
contacts_sum <- contacts_avg <- contacts_lwr <- contacts_upr <-
  matrix(
    data = NA, nrow = length(species), ncol = length(species),
    dimnames = list(species, species)
  )

for (j in species) {
  # Number of hosts j contacted by each (susceptible) host j (within-species)
  counts <-
    apply(
      st_intersects(get(j), sparse = FALSE) &
        outer(get(j)$datetime, get(j)$datetime, "-") >= 0 &
        outer(get(j)$datetime, get(j)$datetime, "-") <=
          time_limit * 24 * 3600,
      1, function(i) {
        ifelse(j %in% c("ibex", "chamois"),
          sum(get(j)$number[i]), sum(i)
        )
      }
    ) - 1 # To avoid counting direct contact with itself

  if (j %in% c("ibex", "chamois")) {
    counts <- rep(counts, times = get(j)$number)
  }

  contacts_sum[j, j] <- sum(counts)
  contacts_avg[j, j] <- 1
  contacts_lwr[j, j] <- 1
  contacts_upr[j, j] <- 1
```

```r
  # Number of hosts k contacted by each (susceptible) host j (between-species)
  for (k in species[species != j]) {
    counts <-
      apply(
        st_intersects(get(j), get(k), sparse = FALSE) &
          outer(get(j)$datetime, get(k)$datetime, "-") >= 0 &
          outer(get(j)$datetime, get(k)$datetime, "-") <=
            time_limit * 24 * 3600,
        1, function(i) {
          ifelse(k %in% c("ibex", "chamois"),
            sum(get(k)$number[i]), sum(i)
          )
        }
      )

    if (j %in% c("ibex", "chamois")) {
      counts <- rep(counts, times = get(j)$number)
    }

    contacts_sum[j, k] <- sum(counts)
    contacts_avg[j, k] <- contacts_sum[j, k] / contacts_sum[j, j]
    contacts_lwr[j, k] <-
      rateratio.test(
        x = c(contacts_sum[j, k], contacts_sum[j, j]),
        n = rep(ifelse(j %in% c("ibex", "chamois"),
          sum(get(j)$number), nrow(get(j))
        ), 2)
      )$conf.int[1]
    contacts_upr[j, k] <-
      rateratio.test(
        x = c(contacts_sum[j, k], contacts_sum[j, j]),
        n = rep(ifelse(j %in% c("ibex", "chamois"),
          sum(get(j)$number), nrow(get(j))
        ), 2)
      )$conf.int[2]
  }
}
```

The estimates of the relative contact rate $\omega_{jk}$ (Figure S1A) were:

```
##            cattle    sheep    goats     ibex  chamois
## cattle  1.000000 0.000000 0.002806 0.015631 0.000000
## sheep   0.000000 1.000000 0.000000 0.082301 0.001271
## goats   1.136364 0.000000 1.000000 3.545455 0.500000
## ibex    0.001465 0.002870 0.001597 1.000000 0.011359
## chamois 0.000000 0.000175 0.000526 0.385141 1.000000
```

The lower bounds of the 95% CI of $\omega_{jk}$ were:

```
##            cattle    sheep    goats     ibex  chamois
## cattle  1.000000 0.000000 0.001533 0.012334 0.000000
## sheep   0.000000 1.000000 0.000000 0.072226 0.000346
## goats   0.614929 0.000000 1.000000 2.185528 0.218925
## ibex    0.001216 0.002517 0.001337 1.000000 0.010643
## chamois 0.000000 0.000004 0.000108 0.366485 1.000000
```

The upper bounds of the 95% CI of $\omega_{jk}$ were:

```
##            cattle    sheep    goats     ibex  chamois
## cattle  1.000000 0.000740 0.004712 0.019546 0.000740
## sheep   0.001173 1.000000 0.001173 0.093446 0.003258
## goats   2.114147 0.182554 1.000000 5.978193 1.075841
## ibex    0.001749 0.003258 0.001893 1.000000 0.012111
## chamois 0.000647 0.000977 0.001537 0.404637 1.000000
```

#### 1.2 Estimates of the force of infection

We estimated the average annual force of infection of each host by minimising the negative log-likelihood for each host species, assuming that the observed number of infected hosts $M_j$ followed a binomial distribution with the number of tested hosts $N_j$ as the number of trials and the prevalence of infection at equilibrium ($i^*$ or $i^* + r^*$) as the probability of success on each trial.

Parameters were estimated by the maximum likelihood method, using the 'optimize' function. The 95% profile likelihood confidence intervals (CIs) were determined by including all the values of the parameter that produced negative log-likelihoods below $-\hat{L} + \frac{\chi^2(0.95)}{2}$, where $-\hat{L}$ was the overall minimum log-likelihood.

Because the prevalence data was pooled for small ruminant herds (no distinction between goat herds and sheep herds), we obtained the same force of infection for both sheep and goats.

```r
# Proportion of infected herds at equilibrium
prop_inf_herds <- function(lambda, gamma) {
  lambda / (lambda + gamma)
}

# Proportion of infected ibex at equilibrium
prop_inf_ibex <- function(lambda, mu) {
  lambda / (lambda + mu)
}

# Proportion of infected chamois at equilibrium
prop_inf_chamois <- function(lambda, gamma, mu) {
  lambda / (lambda + gamma + mu)
}

# Estimate the force of infection and 95% CI derived from profile-likelihood
estim <- function(x, size, species, gamma, mu) {
  estimate <- optimize(function(lambda) {
    -dbinom(
      x,
      size,
      prob =
        if (species == "ibex") {
          prop_inf_ibex(lambda, mu)
        } else if (species == "chamois") {
          prop_inf_chamois(lambda, gamma, mu)
        } else {
          prop_inf_herds(lambda, gamma)
        },
      log = TRUE
    )
  }, interval = c(0, 1))
  profile <- function(lambda) {
    -dbinom(
      x,
      size,
      prob =
        if (species == "ibex") {
          prop_inf_ibex(lambda, mu)
        } else if (species == "chamois") {
          prop_inf_chamois(lambda, gamma, mu)
        } else {
          prop_inf_herds(lambda, gamma)
        },
```

```r
      log = TRUE
    ) - (estimate$objective + 0.5 * qchisq(0.95, 1))
  }
  conf_int <- c(
    ifelse(all(sign(profile(c(0, estimate$minimum))) == -1),
      0, uniroot(profile, interval = c(0, estimate$minimum))$root
    ),
    uniroot(profile, interval = c(estimate$minimum, 1))$root
  )
  return(c(estimate$minimum, conf_int))
}

# Prevalence data
dat <- matrix(
  dat = c(c(1, 0, 0, 32, 1), c(161, 45, 45, 80, 39)),
  nrow = length(species), ncol = 2,
  dimnames = list(species, c("n_pos", "n_test"))
)

# Model parameters
param <- matrix(
  dat = c(c(1, 1, 1, 0.17, 1 / 0.5), c(NA, NA, NA, 0.14, 0.13)),
  nrow = length(species), ncol = 2,
  dimnames = list(species, c("gamma", "mu"))
)

# Initialisation
lambda <- matrix(
  dat = NA, nrow = length(species), ncol = 3,
  dimnames = list(species, c("mean", "lower", "upper"))
)

# Estimation of lambda
for (spec in species) {
  lambda[spec, ] <-
    estim(
      x = dat[spec, "n_pos"],
      size = dat[spec, "n_test"],
      species = spec,
      gamma = param[spec, "gamma"],
      mu = param[spec, "mu"]
    )
}
lambda[lambda[, "mean"] < .Machine$double.eps^0.25, "mean"] <- 0
```

The estimates and 95% CI of the average annual force of infection for each species were:

```
##           mean  lower  upper
## cattle  0.0062 0.0004 0.0278
## sheep   0.0000 0.0000 0.0437
## goats   0.0000 0.0000 0.0437
## ibex    0.0933 0.0592 0.1453
## chamois 0.0561 0.0032 0.2581
```

#### 1.3 Estimates of the transmission rates

The transmission rates were derived from the force of infection, the total number of observations between each pair of hosts, and the prevalence of infection at equilibrium, which were all calculated at previous steps.

The derivation of the corresponding confidence intervals was not straightforward as the transmission rates $\beta$ combine several parameter estimates. Therefore, we simulated 100 000 values of the prevalence of infection and of the number of observed contacts, assuming their distribution was $Beta(M_j + 0.5, N_j - M_j + 0.5)$ and $Gamma((\sum n_{jk}) + 0.5, 1)$, respectively, which is equivalent to using the Jeffreys interval for a binomial proportion or a Poisson rate.

```
n_sim <- 100000

# Force of infection in herds at equilibrium
lambda_herds <- function(p, gamma) {
  gamma * p / (1 - p)
}

# Force of infection in ibex at equilibrium
lambda_ibex <- function(p, mu) {
  mu * p / (1 - p)
}

# Force of infection in chamois at equilibrium
lambda_chamois <- function(p, gamma, mu) {
  (gamma + mu) * p / (1 - p)
}

# Initialisation
pinf_sim <- lambda_sim <- matrix(NA,
  nrow = n_sim, ncol = length(species),
  dimnames = list(NULL, species)
)
```

```r
# Simulations of lambda
for (j in species) {
  pinf_sim[, j] <-
    rbeta(
      n_sim,
      dat[j, "n_pos"] + 0.5,
      dat[j, "n_test"] - dat[j, "n_pos"] + 0.5
    )

  lambda_sim[, j] <-
    if (j == "ibex") {
      lambda_ibex(
        pinf_sim[, j],
        param[j, "mu"]
      )
    } else if (j == "chamois") {
      lambda_chamois(
        pinf_sim[, j],
        param[j, "gamma"],
        param[j, "mu"]
      )
    } else {
      lambda_herds(
        pinf_sim[, j],
        param[j, "gamma"]
      )
    }
}

# Simulations of contacts
contacts_sim <-
  sapply(contacts_sum, function(x) rgamma(n_sim, shape = x + 0.5, rate = 1))
```

For each simulation, we calculated $\lambda_{j0}$, $\Lambda_{jk}$ and $i_k^*$ and the corresponding $\beta_{jk}$. The credible interval (CrI) was then calculated as the equal-tailed 95% interval of the distribution of the 100 000 values of $\beta_{jk}$. This allowed us to obtain each element $\beta_{jk}$ of the 'who acquires infection from whom' (WAIFW) matrix.

```r
# Initialisation
waifw_avg <- waifw_lwr <- waifw_upr <-
  matrix(
    data = NA, nrow = length(species), ncol = length(species),
    dimnames = list(species, species)
  )
```

```r
beta <- matrix(data = NA, nrow = n_sim, ncol = length(species)^2)

# Proportion of infectious ibex at equilibrium
prop_inf_ibex2 <- function(lambda, gamma, mu) {
  (lambda * mu) / ((gamma + mu) * (lambda + mu))
}

# Within-species transmission rate
beta_intra <- function(par) {
  par[j] * par[length(species) + j] / (
    prop_inf_herds(par[1], param[1, "gamma"]) * par[6] +
      prop_inf_herds(par[2], param[2, "gamma"]) * par[7] +
      prop_inf_herds(par[3], param[3, "gamma"]) * par[8] +
      prop_inf_ibex2(par[4], param[4, "gamma"], param[4, "mu"]) * par[9] +
      prop_inf_chamois(par[5], param[5, "gamma"], param[5, "mu"]) * par[10])
}

# Between-species transmission rate
beta_inter <- function(par) {
  par[j] * par[length(species) + k] / (
    prop_inf_herds(par[1], param[1, "gamma"]) * par[6] +
      prop_inf_herds(par[2], param[2, "gamma"]) * par[7] +
      prop_inf_herds(par[3], param[3, "gamma"]) * par[8] +
      prop_inf_ibex2(par[4], param[4, "gamma"], param[4, "mu"]) * par[9] +
      prop_inf_chamois(par[5], param[5, "gamma"], param[5, "mu"]) * par[10])
}

for (j in seq_len(length(species))) {
  # Transmission rate within species j
  waifw_avg[j, j] <-
    beta_intra(par = c(lambda[, "mean"], contacts_avg[j, ]))

  tmp <-
    apply(
      cbind(lambda_sim, contacts_sim[, j + seq(0, 24, 5)]),
      1, function(i) beta_intra(i)
    )
  beta[, j + seq(0, 24, 5)[j]] <- tmp
  waifw_lwr[j, j] <- quantile(tmp, 0.025)
  waifw_upr[j, j] <- quantile(tmp, 0.975)

  for (k in seq_len(length(species))[-j]) {
    # Transmission rate from species k to species j
    waifw_avg[j, k] <-
```

```r
      beta_inter(par = c(lambda[, "mean"], contacts_avg[j, ]))

    tmp <-
      apply(
        cbind(lambda_sim, contacts_sim[, j + seq(0, 24, 5)]),
        1, function(i) beta_inter(i)
      )
    beta[, k + seq(0, 24, 5)[j]] <- tmp
    waifw_lwr[j, k] <- quantile(tmp, 0.025)
    waifw_upr[j, k] <- quantile(tmp, 0.975)
  }
}
waifw_lwr[waifw_avg == 0] <- 0
```

The estimates of the transmission rates $\beta_{jk}$ (Figure S2A) were:

```
##           cattle  sheep  goats   ibex chamois
## cattle    0.6911 0.0000 0.0019 0.0108  0.0000
## sheep     0.0000 0.0000 0.0000 0.0000  0.0000
## goats     0.0000 0.0000 0.0000 0.0000  0.0000
## ibex      0.0008 0.0015 0.0008 0.5158  0.0059
## chamois   0.0000 0.0001 0.0003 0.2267  0.5887
```

The lower bounds of the 95% CrI of $\beta_{jk}$ were:

```
##           cattle  sheep goats   ibex chamois
## cattle    0.1890 0.0000 5e-04 0.0030  0.0000
## sheep     0.0000 0.0000 0e+00 0.0000  0.0000
## goats     0.0000 0.0000 0e+00 0.0000  0.0000
## ibex      0.0006 0.0012 7e-04 0.4397  0.0049
## chamois   0.0000 0.0000 0e+00 0.0317  0.0823
```

The upper bounds of the 95% CrI of $\beta_{jk}$ were:

```
##           cattle  sheep  goats   ibex chamois
## cattle    0.9402 0.0004 0.0036 0.0159  0.0004
## sheep     0.0003 0.8318 0.0003 0.0688  0.0016
## goats     0.0964 0.0029 0.0847 0.2850  0.0440
## ibex      0.0010 0.0019 0.0011 0.6303  0.0072
## chamois   0.0004 0.0007 0.0014 0.5835  1.5113
```

#### 1.4 Next-generation matrix

Finally, the next-generation matrix $K$ is given by $K = -T\Sigma^{-1}$, where $\Sigma$ is the transition matrix, i.e. the $n \times n$ matrix corresponding to recovery and death of state $i$. Therefore, $-\Sigma^{-1}$ describes the average duration that an individual/herd will stay in the infectious state $i$. Each element of $K$ can be interpreted in a similar way as the basic reproduction number: $K_{jk}$ is the expected number of secondary cases in host $j$ that would arise from the introduction of one infectious host $k$ in a susceptible population of host $j$. The overall basic reproduction number $R_0$ in the multi-host system is the dominant eigenvalue of $K$.

We also calculated the host-specific reproduction number $U_m = \rho(P_m K)$ (where $P_m$ is the $n \times n$ projection matrix), measuring the expected number of secondary infections in hosts $m$ resulting from an average infectious individual of hosts $m$ in the absence of other hosts; and the host-excluded reproduction number $Q_m = \rho((I - P_m)K)$ (where $I$ is the $n \times n$ identity matrix), measuring the expected number of secondary infections in hosts other than $m$ resulting from an average infectious individual of hosts other than $m$ in the absence of hosts $m$.

```r
# Transition matrix
sigma <-
  diag(
    x = c(
      -param["cattle", "gamma"],
      -param["sheep", "gamma"],
      -param["goats", "gamma"],
      -(param["ibex", "gamma"] + param["ibex", "mu"]),
      -(param["chamois", "gamma"] + param["chamois", "mu"])
    ),
    nrow = length(species), ncol = length(species)
  )
colnames(sigma) <- rownames(sigma) <- species

# Next-generation matrix
ngm_avg <- waifw_avg %*% (-solve(sigma))
eigenvalues_avg <- eigen(ngm_avg)$values
r0_avg <- max(Re(eigenvalues_avg[abs(Im(eigenvalues_avg)) < 1e-6])) # R0

# Lower bound of the 95% credible interval
ngm_lwr <- waifw_lwr %*% (-solve(sigma))
eigenvalues_lwr <- eigen(ngm_lwr)$values
r0_lwr <- max(Re(eigenvalues_lwr[abs(Im(eigenvalues_lwr)) < 1e-6]))

# Upper bound of the 95% credible interval
ngm_upr <- waifw_upr %*% (-solve(sigma))
eigenvalues_upr <- eigen(ngm_upr)$values
r0_upr <- max(Re(eigenvalues_upr[abs(Im(eigenvalues_upr)) < 1e-6]))
```

The estimates of $K_{jk}$ (Figure S3A) were:

```
##          cattle  sheep  goats   ibex chamois
## cattle  0.6911 0.0000 0.0019 0.0348  0.0000
## sheep   0.0000 0.0000 0.0000 0.0000  0.0000
## goats   0.0000 0.0000 0.0000 0.0000  0.0000
## ibex    0.0008 0.0015 0.0008 1.6640  0.0028
## chamois 0.0000 0.0001 0.0003 0.7313  0.2764
```

The lower bounds of the 95% CrI of $K_{jk}$ were:

```
##          cattle  sheep goats   ibex chamois
## cattle  0.1890 0.0000 5e-04 0.0096  0.0000
## sheep   0.0000 0.0000 0e+00 0.0000  0.0000
## goats   0.0000 0.0000 0e+00 0.0000  0.0000
## ibex    0.0006 0.0012 7e-04 1.4185  0.0023
## chamois 0.0000 0.0000 0e+00 0.1023  0.0386
```

The upper bounds of the 95% CrI of $K_{jk}$ were:

```
##          cattle  sheep  goats   ibex chamois
## cattle  0.9402 0.0004 0.0036 0.0512  0.0002
## sheep   0.0003 0.8318 0.0003 0.2220  0.0008
## goats   0.0964 0.0029 0.0847 0.9192  0.0207
## ibex    0.0010 0.0019 0.0011 2.0332  0.0034
## chamois 0.0004 0.0007 0.0014 1.8823  0.7096
```

The overall $R_0$ was 1.67 (95% CrI: 1.42, 2.04).

```r
# Host-specific reproduction number U
p <- matrix(
  data = 0, nrow = length(species), ncol = length(species),
  dimnames = list(species, species)
)
p["ibex", "ibex"] <- 1 # projection matrix

eigenvalues_avg <- eigen(p %*% ngm_avg)$values
u_avg <- max(Re(eigenvalues_avg[abs(Im(eigenvalues_avg)) < 1e-6]))
eigenvalues_lwr <- eigen(p %*% ngm_lwr)$values
u_lwr <- max(Re(eigenvalues_lwr[abs(Im(eigenvalues_lwr)) < 1e-6]))
eigenvalues_upr <- eigen(p %*% ngm_upr)$values
u_upr <- max(Re(eigenvalues_upr[abs(Im(eigenvalues_upr)) < 1e-6]))
```

```r
# Host-excluded reproduction number Q
i <- diag(length(species)) # identity matrix

eigenvalues_avg <- eigen((i - p) %*% ngm_avg)$values
q_avg <- max(Re(eigenvalues_avg[abs(Im(eigenvalues_avg)) < 1e-6]))
eigenvalues_lwr <- eigen((i - p) %*% ngm_lwr)$values
q_lwr <- max(Re(eigenvalues_lwr[abs(Im(eigenvalues_lwr)) < 1e-6]))
eigenvalues_upr <- eigen((i - p) %*% ngm_upr)$values
q_upr <- max(Re(eigenvalues_upr[abs(Im(eigenvalues_upr)) < 1e-6]))
```

The ibex-specific reproduction number $U$ was 1.66 (95% CrI: 1.42, 2.03). The ibex-excluded reproduction number $Q$ was 0.69 (95% CrI: 0.19, 0.94).

### 2 Sensitivity analysis

We assessed the sensitivity of our results to two critical model assumptions: the survival time of *Brucella* in pastures, used to estimate the relative contact rates, and the disease-induced mortality in chamois (see Figures S1B-D, S2B-D, S3B-D).

#### 2.1 Survival time in the environment

For the survival time of *Brucella* in the mountain environment, we evaluated two values in addition to the baseline value of 25 days: a lower value of 15 days and a higher value taking into account all possible indirect contacts during the observation period (maximum of 49 days).

Assuming a survival time of 15 days instead of 25 days, the overall $R_0$ was 1.67 (95% CrI: 1.42, 2.04). The ibex-specific reproduction number $U$ was 1.66 (95% CrI: 1.42, 2.03). The ibex-excluded reproduction number $Q$ was 0.65 (95% CrI: 0.16, 0.93). The estimates of $K_{jk}$ (Figure S3B) were:

```
##         cattle  sheep  goats   ibex chamois
## cattle  0.6537 0.0000 0.0017 0.0390  0.0000
## sheep   0.0000 0.0000 0.0000 0.0000  0.0000
## goats   0.0000 0.0000 0.0000 0.0000  0.0000
## ibex    0.0008 0.0016 0.0004 1.6639  0.0028
## chamois 0.0000 0.0001 0.0001 0.7168  0.2912
```

The lower bounds of the 95% CrI of $K_{jk}$ were:

```
##         cattle  sheep goats   ibex chamois
## cattle  0.1648 0.0000 4e-04 0.0099  0.0000
## sheep   0.0000 0.0000 0e+00 0.0000  0.0000
## goats   0.0000 0.0000 0e+00 0.0000  0.0000
## ibex    0.0006 0.0013 3e-04 1.4184  0.0024
## chamois 0.0000 0.0000 0e+00 0.1020  0.0415
```

The upper bounds of the 95% CrI of $K_{jk}$ were:

```
##         cattle  sheep  goats   ibex chamois
## cattle  0.9263 0.0004 0.0034 0.0590  0.0002
## sheep   0.0005 0.8622 0.0005 0.1941  0.0011
## goats   0.1158 0.0035 0.0924 0.9294  0.0062
## ibex    0.0011 0.0020 0.0006 2.0332  0.0035
## chamois 0.0004 0.0008 0.0008 1.7960  0.7285
```

Assuming a survival time of 49 days instead of 25 days, the overall $R_0$ was 1.67 (95% CrI: 1.42, 2.04). The ibex-specific reproduction number $U$ was 1.66 (95% CrI: 1.42, 2.03). The ibex-excluded reproduction number $Q$ was 0.66 (95% CrI: 0.16, 0.93). The estimates of $K_{jk}$ (Figure S3C) were:

```
##         cattle  sheep  goats   ibex chamois
## cattle  0.6556 0.0000 0.0074 0.0388  0.0000
## sheep   0.0000 0.0000 0.0000 0.0000  0.0000
## goats   0.0000 0.0000 0.0000 0.0000  0.0000
## ibex    0.0007 0.0014 0.0008 1.6640  0.0027
## chamois 0.0000 0.0001 0.0003 0.7472  0.2601
```

The lower bounds of the 95% CrI of $K_{jk}$ were:

```
##         cattle  sheep  goats   ibex chamois
## cattle  0.1621 0.0000 0.0018 0.0097  0.0000
## sheep   0.0000 0.0000 0.0000 0.0000  0.0000
## goats   0.0000 0.0000 0.0000 0.0000  0.0000
## ibex    0.0006 0.0011 0.0006 1.4186  0.0023
## chamois 0.0000 0.0000 0.0000 0.1025  0.0357
```

The upper bounds of the 95% CrI of $K_{jk}$ were:

```
##         cattle  sheep  goats   ibex chamois
## cattle  0.9245 0.0003 0.0117 0.0580  0.0002
## sheep   0.0003 0.7512 0.0003 0.3015  0.0006
## goats   0.0843 0.0023 0.0932 0.9164  0.0169
## ibex    0.0009 0.0018 0.0010 2.0333  0.0034
## chamois 0.0003 0.0007 0.0013 1.9768  0.6885
```

#### 2.2 SIR model in chamois

The assumption that all infectious chamois died was based on past observations of a few sporadic cases of *B. abortus* and *B. melitensis* in chamois in Europe, which displayed very severe lesions suggesting disease-induced mortality. High disease-induced mortality in chamois could also explain why self-sustained persistence of *Brucella* in the absence of transmission from domestic species have never been observed in this species. However, some individual chamois do not seem to display symptoms. In the study area, there were only a few cases, including a single case in found-dead animals. Therefore, the information was scarce in chamois compared to the available knowledge in ibex, which led us to evaluate an alternative SIR model for chamois. In the absence of specific data, we used the same recovery rate in chamois than in ibex (0.17 per year). The reality is probably in-between these two model formulations, but further data would be required to gain a better understanding of recovery and brucellosis-induced mortality in chamois.

When using a SIR model in chamois, the overall $R_0$ was 1.67 (95% CrI: 1.42, 2.04). The ibex-specific reproduction number $U$ was 1.67 (95% CrI: 1.42, 2.04). The ibex-excluded reproduction number $Q$ was 0.69 (95% CrI: 0.19, 0.94). The estimates of $K_{jk}$ (Figure S3D) were:

```
##         cattle  sheep  goats   ibex chamois
## cattle  0.6911 0.0000 0.0019 0.0348  0.0000
## sheep   0.0000 0.0000 0.0000 0.0000  0.0000
## goats   0.0000 0.0000 0.0000 0.0000  0.0000
## ibex    0.0008 0.0015 0.0008 1.6655  0.0195
## chamois 0.0000 0.0000 0.0000 0.0526  0.1411
```

The lower bounds of the 95% CrI of $K_{jk}$ were:

```
##         cattle  sheep goats   ibex chamois
## cattle  0.1884 0.0000 5e-04 0.0094  0.0000
## sheep   0.0000 0.0000 0e+00 0.0000  0.0000
## goats   0.0000 0.0000 0e+00 0.0000  0.0000
## ibex    0.0006 0.0012 7e-04 1.4217  0.0165
## chamois 0.0000 0.0000 0e+00 0.0063  0.0171
```

The upper bounds of the 95% CrI of $K_{jk}$ were:

```
##         cattle  sheep  goats   ibex chamois
## cattle  0.9399 0.0004 0.0036 0.0512  0.0012
## sheep   0.0003 0.8327 0.0003 0.2218  0.0053
## goats   0.0978 0.0029 0.0863 0.9358  0.1514
## ibex    0.0010 0.0019 0.0011 2.0356  0.0241
## chamois 0.0000 0.0001 0.0001 0.1777  0.4765
```

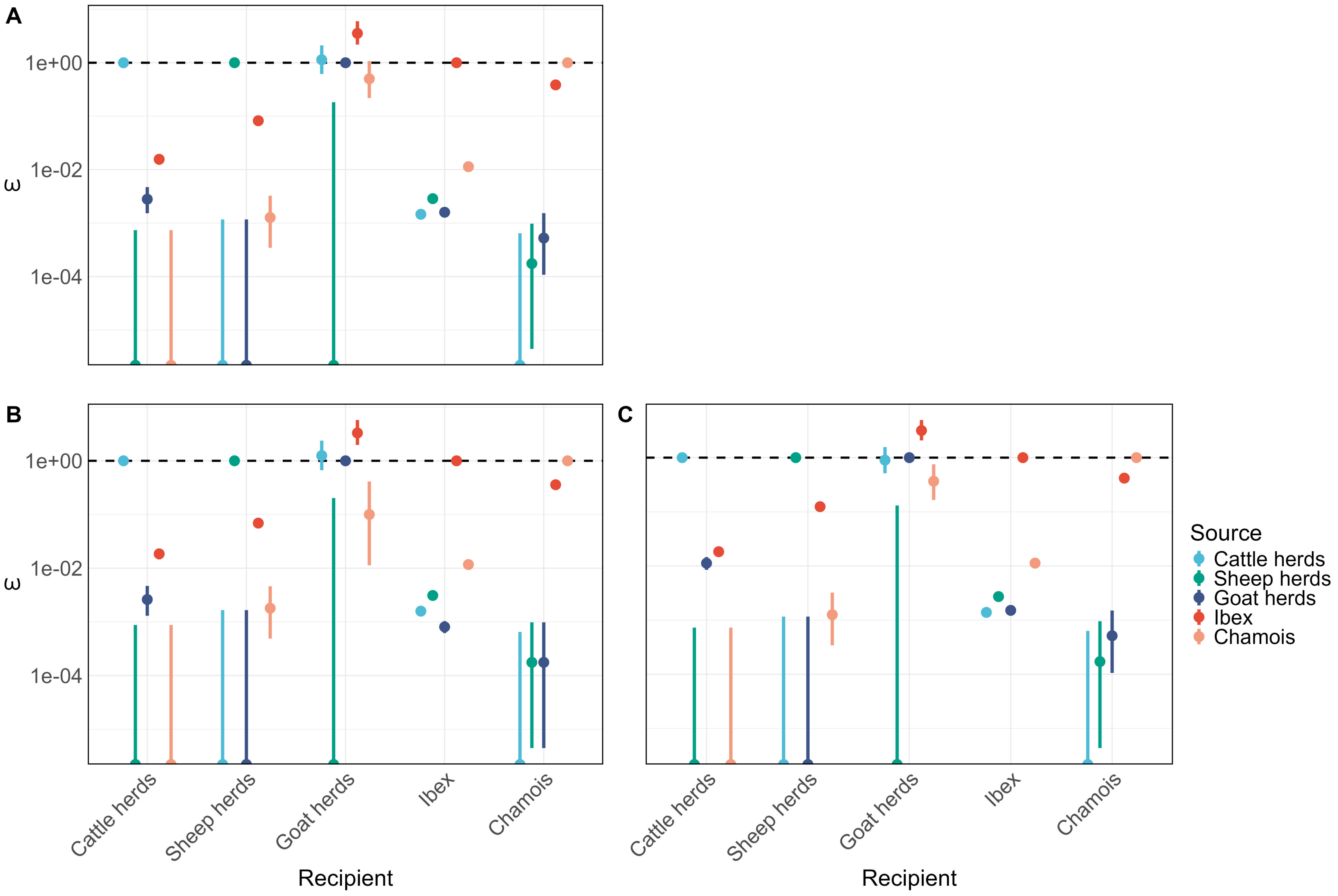

Figure S1: Estimates of the relative contact rates. A: baseline model; B: results for an alternative value of Brucella survival in mountain pastures of 15 days; C: results for an alternative value of Brucella survival in mountain pastures of 49 days. The horizontal dashed line represents the set value for the within-species relative contact rates, set to 1. Note that the y-axis is on the log scale.

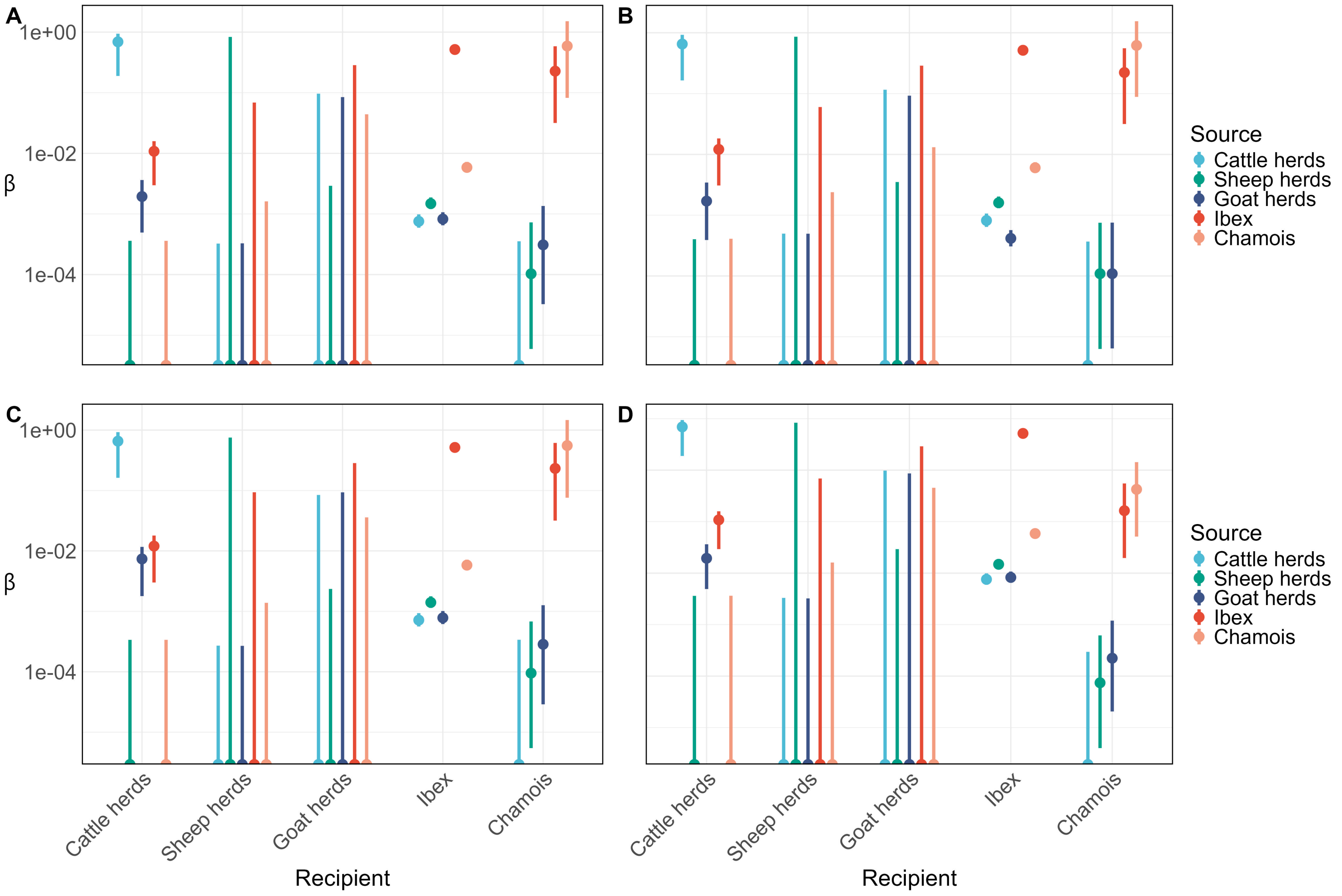

Figure S2: Estimates of the within- and between-species contact rates in the who acquires infection from whom matrix. A: baseline model; B: results for an alternative value of Brucella survival in mountain pastures of 15 days; C: results for an alternative value of Brucella survival in mountain pastures of 49 days; D: results when using a SIR model for chamois instead of a SI model. Note that the y-axis is on the log scale.

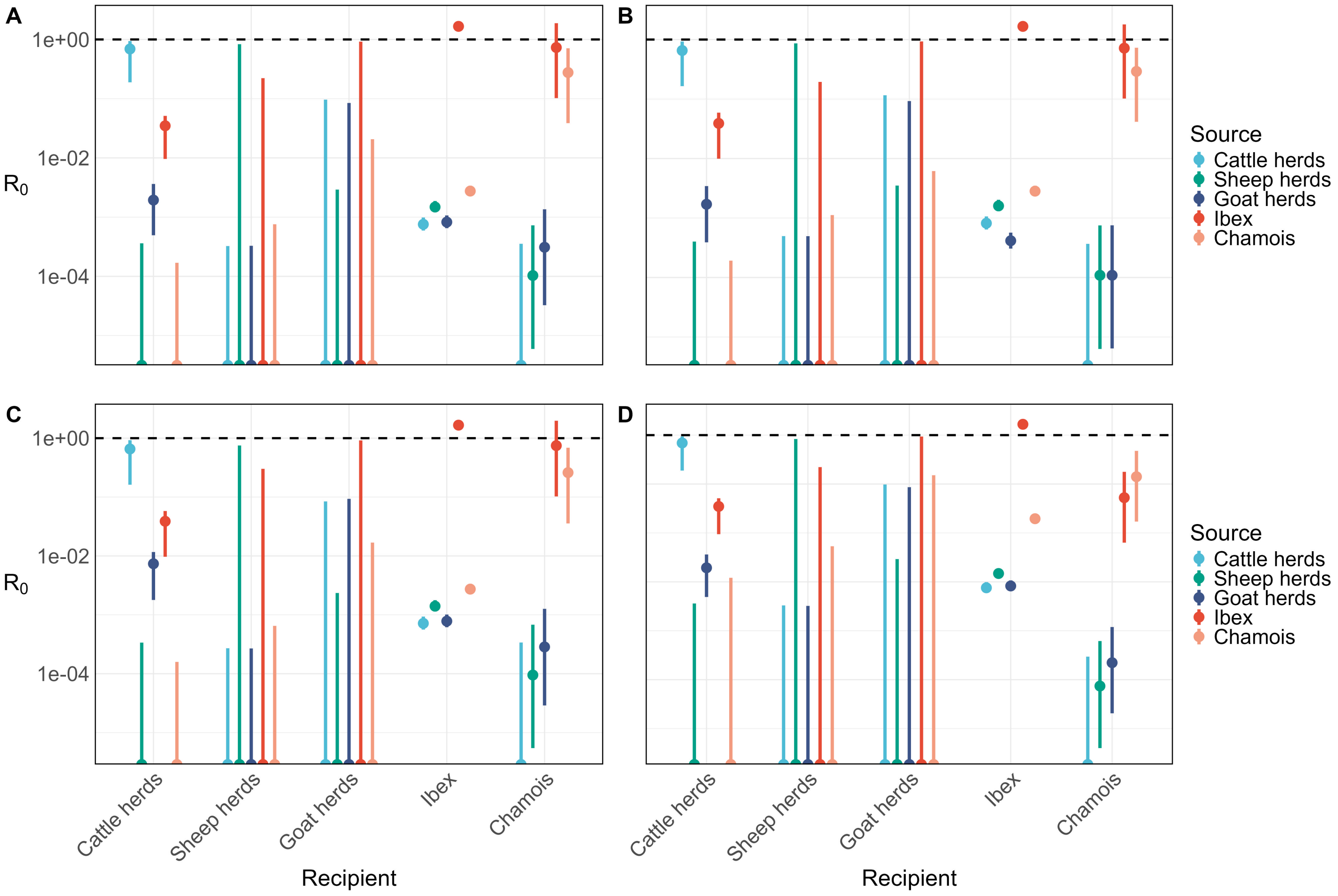

Figure S3: Estimates of the within- and between-species basic reproduction number in the next-generation matrix. A: baseline model; B: results for an alternative value of Brucella survival in mountain pastures of 15 days; C: results for an alternative value of Brucella survival in mountain pastures of 49 days; D: results when using a SIR model for chamois instead of a SI model. The horizontal dashed line represents the threshold for pathogen maintenance. Note that the y-axis is on the log scale.
